# Improving whole biodiversity monitoring and discovery with environmental DNA metagenomics

**DOI:** 10.1101/2024.12.17.628643

**Authors:** Manuel Curto, Ana Veríssimo, Giulia Riccioni, Carlos D. Santos, Filipe Ribeiro, Sissel Jentoft, Maria Judite Alves, Hugo F. Gante

## Abstract

Environmental DNA (eDNA) metagenomics sequences all DNA molecules present in environmental samples and has the potential of identifying virtually any organism from which they are derived. However, due to unacceptable levels of false positives and negatives, this approach is underexplored as a tool for biodiversity monitoring across the tree of life, particularly for non-microscopic eukaryotes. We present SEQIDIST, a framework that combines multilocus BLAST matches against several reference databases followed by analysis of sequence identity distribution patterns to disentangle false positives while revealing new biodiversity and increasing the accuracy of metagenomic approaches. We tested SEQIDIST on an eDNA metagenomic dataset from a riverine site and compare the results to those obtained with an eDNA metabarcoding approach for benchmarking purposes. We start by characterizing the biological community (∼ 2000 taxa) across the tree of life at low taxonomic levels and show that eDNA metagenomics has a higher sensitivity than eDNA metabarcoding in discovering new diversity. We show that limited representation of whole genome sequences in reference databases can lead to false positives. For non-microscopic eukaryotes, eDNA metagenomic data often consist of a few sparse, anonymous sequences scattered across the genome, making metagenome assembly methods unfeasible. Finally, we infer eDNA source and residency time using read length distributions as a measure of decay status. The higher accuracy of SEQIDIST opens the discussion of the archival potential of eDNA metagenomics and its implementation in biodiversity monitoring actions at large planetary and temporal scales.

## 1 Introduction

Global declines and changes in biodiversity distribution are hallmarks of the Anthropocene (Butchart et al., 2010). Moreover, lack of taxonomic expertise, and financial and technical challenges hinder accurate large-scale biodiversity assessment (Pereira et al., 2010). Importantly, much biodiversity is still undocumented, and knowledge gaps are more pronounced in underexplored regions, especially in diverse freshwater and marine environments, where biodiversity loss is likely underestimated (Schmeller et al., 2017).

The DNA present in the environment (eDNA), derived from both living microorganisms (intracellular eDNA) and remains of macroscopic eukaryotes (extracellular eDNA), is a powerful source of biodiversity information (Taberlet et al., 2012). In the context of biodiversity monitoring, eDNA appears as an emergent alternative to observation-based methods that are often invasive, require taxonomic expertise and are inefficient in detecting rare or elusive species (Goldberg et al., 2016). Moreover, eDNA samples harbour information across the tree of life widening their applicability to multispecies monitoring projects (Stat et al., 2017). In this context, rivers are a particularly interesting source of biodiversity information since they accumulate eDNA molecules originating not only from local aquatic organisms but also from upstream and nearby terrestrial biota (Deiner et al., 2016), through long-distance transport, runoff, and possibly also deposition of airborne eDNA (Clare et al., 2022; Littlefair et al., 2023; Lynggaard et al., 2022). Therefore, riverine eDNA samples have the potential to provide biodiversity information at the landscape level.

The analysis of eDNA samples by amplifying and sequencing specific barcodes, known as eDNA metabarcoding, has revolutionized biodiversity monitoring due to its relatively simple and cost-effective implementation (Deiner et al., 2017). Nevertheless, this method has known limitations that need to be circumvented, such as biases in taxon amplification and decay rate (Jo et al., 2017; Kelly et al., 2019). In turn, environmental metagenomics (or shotgun metagenomics (Quince et al., 2017)) sequences DNA molecules from an environmental sample (e.g., water, soil, air) without any taxonomic biases or prior knowledge of species composition. By doing so, eDNA metagenomics is the methodology that can unlock the largest amount of information from eDNA samples.

Despite its proven success in characterizing microbial communities for two decades (Venter et al., 2004), application of eDNA metagenomics to non-microscopic eukaryotes has been surprisingly rare, and severely constrained by analytical limitations leading to high prevalence of false positives and negatives (Simon et al., 2019; Stat et al., 2017). eDNA from non-microscopic eukaryotes is usually extracellular, fragmented, and present in the environment in low abundance (Nagler et al., 2022), making the assembly-based methodologies, already established in the field of microbiology, ineffective in reconstructing larger contigs (Quince et al., 2017). Thus, the detection of from non-microscopic eukaryotes has to rely on directly comparing anonymous short reads to reference databases (Manu & Umapathy, 2023; Singer et al., 2020). Because reference sequence databases are incomplete in terms of taxonomic representation, most of the data remains unmapped resulting in false negatives. Furthermore, some regions of the genome are conserved across species boundaries attracting heterospecific matches that may raise false positives (Simon et al., 2019). The rare studies describing multicellular eukaryotic communities using eDNA metagenomics were conducted at high taxonomic levels, and taxon detection was based on individual sequence matches (Cowart et al., 2018; Manu & Umapathy, 2023; Singer et al., 2020). Here, we address the lack of resolution and detection bias of using single sequence matches by developing a new framework that makes eDNA metagenomics a viable and powerful tool for whole biodiversity monitoring including for non-microscopic organisms.

In this study, we propose using eDNA metagenomics coupled with a dedicated bioinformatics workflow, named SEQIDIST. Instead of considering individual sequence matches as indicators of positive detection, which can lead to false positives, SEQIDIST combines percent identity information from all BLAST matches to a taxon. This approach assumes that while some regions of the genome may be conserved across species boundaries, this will not be the case for most of the genome. Therefore, sequence identity distribution patterns should differ between intraspecific and interspecific comparisons. We provide proof-of-principle for SEQIDIST’s ability to detect false positives. By applying SEQIDIST to an eDNA metagenomic dataset from the Ave River (NW Iberian Peninsula), we demonstrate that SeqIDist can characterize freshwater communities across the tree of life with high accuracy and greater sensitivity than traditional eDNA metabarcoding. In this context, we focus on fish taxa as a key case-study to evaluate the ability of metagenomics to accurately describe eDNA diversity at the species level. Teleost fishes are the best documented taxon in the Ave River and the group of organisms with the larger amount of genomic resources in public databases. Additionally, we test if read length contains information on eDNA decay, which could improve the characterization of eDNA origin (e.g., intra-versus extracellular, and habitat), illustrating the potential of metagenomics to extract additional levels of information from eDNA samples.

## 2 Material and Methods

### 2.1 Multilocus SEQIDIST detection of true and false positives

Most genomic reads are expected to match genome assemblies of conspecific taxa at high sequence identity, with no or very few mismatches. Nevertheless, some proportion of reads can also match heterospecific taxa with high sequence identity (e.g., >99%), which would result in false positives if individual matches of each read are considered. This can happen because of missing representative genomes in the reference database, incomplete assemblies, or high degree of conservation of some genomic regions. Nevertheless, only a small part of the genome should be conserved across species making heterospecific matches with high sequence identity much less frequent than heterospecific matches with lower sequence identity. Therefore, we expect that the multilocus sequence identity distribution of metagenomics reads to a certain taxon can provide information on whether its detection is a true or false positive. Specifically, false positives should show proportionally more multilocus matches with lower identity values. We tested this expectation by evaluating the sequence identity distributions of BLAST hits from shotgun sequence data against genomes assemblies across a range of intraspecific, intrageneric, and intrafamilial comparisons. Shotgun Illumina sequencing datasets without any type of enrichment were selected for a few case-study fish species likely to be detected in our study area (described in section 2.2 below). These included at least one sample of species represented in the short-read archive (SRA) database from GenBank for the following genera (Table S1): *Gobio*, *Gambusia*, *Lepomis*, *Micropterus*, *Luciobarbus*, *Perca*, *Sander*, *Salmo*, and *Squalius*.

Short reads were downloaded with the program fastq-dump from the sra toolkit v. 3.0.0 (Sequence Read Archive Toolkit) (https://www.ncbi.nlm.nih.gov/sra/docs/toolkitsoft), and filtered for quality with trimmomatic (Bolger et al., 2014) (details in *2.5. Quality control of shotgun reads*). Possible PCR duplicates were removed with the script clumpify.sh from BBMap (Bushnell, 2014). To mimic the low coverage data obtained with metagenomics data, 1000 random reads were selected from the dataset with seqtk (Li, 2012). Paired reads were then merged with PEAR (Zhang et al., 2014) with default parameters and converted to fasta format. Reads were then compared to individual reference genomes (Table S1) using megablast searches and matches were processed as described in the section *2.7*. *Taxonomic assignment of metagenomics reads*. The output is a table with the BLAST match parameters per read and the corresponding taxonomic assignment. These were used in R (R Core Team, 2023) to produce histograms based on match sequence identity using the library ggplot2 (Wickham et al., 2016). Several bin sizes were tested (binwidth = 0.1, 0.5, 1.0, 1.25), with binwidth parameter of 1.25 providing maximum differentiation between intra- and interspecific comparisons, although the overall pattern did not change for other values. The bin encompassing the highest sequence identity included matches of ID >99.5%. The remaining bins corresponded to intervals of 1% sequence identity.

### 2.2 Study area and sampling site

The Ave River is a small drainage located in northwestern Iberian Peninsula, flowing for about 91 km, from the Cabreira mountain to the Atlantic Ocean at Vila do Conde, Portugal (in a North-East to South-West direction). This drainage (41°15–40°45N; 7°45–8°45W) encompasses an area of 1473 km^2^, ranging up to 1261 meters altitude. The climate is mildly Mediterranean (Koppen-Geiger classification, Csb) with cool summers, and strong Atlantic influence. Mean annual rainfall is about 1400 mm (ranging from 900–3900 mm) and the annual mean temperature of 12–15° C. The basin drains mostly from limestone rocks, with high groundwater input, and land cover is mostly forest (57%, mainly pine and eucalyptus) and farmland (34%). However, the lower and middle sections of the drainage are heavily modified and degraded due to heavy industrialization and urbanization. The area around this drainage is inhabited by more than 400 people per km^2^, being more densely inhabited in the middle and lower sections. Despite the significant infrastructure efforts in water treatment, the Ave River basin is still considered one of the most polluted rivers in Portugal; yet, the upper Ave watershed and main tributaries are generally pristine areas, populated mostly by brown trout (*Salmo trutta*) and native leuciscids (Collares-Pereira et al., 2021). In contrast, the lower and middle sections of the Ave watershed are generally dominated by non-native fish species, with a small number of migratory species, such as European eel (*Anguilla anguilla*) and thinlip grey mullet (*Chelon ramada*) (Collares-Pereira et al., 2021).

Water samples were collected in the lower section of the Ave River (41°21’6.93"N; 8°40’54.83"W) at ∼9.5 km from the river mouth, and just downstream from its fourth barrier (an abandoned watermill). Downstream barriers are semi-permeable and in high winter river flow conditions allow fishes to move upstream. Fish communities in this region are composed of the Pyrenean gudgeon (*Gobio lozanoi*), pumpkinseed sunfish (*Lepomis gibbosus*), Iberian barbel (*Luciobarbus bocagei*), European eel, eastern mosquitofish (*Gambusia holbrooki*), northern Iberian chub (*Squalius carolitertii*), northern straight-mouth nase (*Pseudochondrostoma duriense*), ruivaco (*Achondrostoma oligolepis*), thinlip grey mullet and roach (*Rutilus rutilus*) (Collares-Pereira et al., 2021; Ribeiro & Veríssimo, 2014). Although migratory species such as *Alosa* spp. and *Salmo trutta* may occur, their presence is unlikely since their migration is highly constrained by the barriers present downstream of the sampling site.

### 2.3 Sampling and DNA isolation

Five 2 L water samples were collected on site using autoclaved water bottles, transported on ice and in the dark to the laboratory, and filtered within 6 h upon collection. Filtration was performed using high-capacity nitrocellulose filters (700 cm^2^ filter area) with 0.45 µm pore diameter (GoPro^TM^, Proactive environmental products, Lakewood Ranch, Florida, USA), attached to sterile tubing and a peristaltic Millipore vacuum pump. Upon filtration, the filters were filled with a 40 mL solution of lysis buffer and lysis additive (ratio of 3:1), following the protocol of (Sellers et al., 2018), and sealed with sterile rubber stoppers. All filters were kept in the fridge until eDNA extraction (up to 5 months). Details on DNA isolation are available in supplementary methods.

### 2.4 Shotgun library preparation and sequencing

Library preparation and sequencing was performed by the Norwegian Sequencing Centre (www.sequencing.uio.no), University of Oslo, Norway. Library preparation followed the SMARTer ThruPLEX® (Takara Bio, Kusatsu, Shiga, Japan) low-input sample prep protocol, including sonication of samples on a Covaris LE220 aiming at ∼350 bp fragments. The final pooled libraries were sequenced using two runs of the Illumina MiSeq v2 500 cycles aiming at a total of 4 million reads per extraction (all samples run in both runs).

### 2.5 Quality control of shotgun reads

Shotgun sequencing reads were quality-trimmed with Trimmomatic (Bolger et al., 2014) by excluding adapters at the 3’-end of the reads, and regions with quality below 20. The resulting paired reads were merged using PEAR (Zhang et al., 2014) with default parameters. For further analyses, both merged and unmerged reads longer than 50 bp were considered.

### 2.6 Metagenomics reference databases

Taxonomic assignment was done against four reference databases (Table S1): *a*) all non-redundant *nucleotide* sequences from GenBank (accessed in December 2020); *b*) all transcriptomes of fish families present in Portuguese freshwater bodies available in GenBank (*transcriptomes*, accessed in May 2021) and detected in the *nt* database; *c*) whole genome sequences of species of fish families found in Portuguese freshwater bodies available in GenBank and detected in the *nt* database (*genomes_v1*; accessed in May 2021); *d*) an extended fish genome database including the most recent whole genome assemblies of species in fish families detected with databases *a*) to *c*) (*genomes_v2*; accessed in July 2022), plus newly generated whole genome sequence data from fish taxa present in Portuguese river drainages (Table S1). The *nt* database was chosen to allow taxonomic assignment of reads across the tree of life at different taxonomic levels. Databases *b*) to *d*) focused on the identification of fish species only to evaluate database requirements for classification at the species level. During data analysis and writing of this manuscript, several new fish genomes were produced by our team and others became available on GenBank, filling some of the gaps existing in the *genomes_v1* database *c*). Thus, the extended *genomes_v2* database was used to evaluate how completeness of genomic reference databases might affect detections at the species level. In summary, the *transcriptomes* database *b*) is composed of 233 transcriptomes from 28 fish families and 120 species. The *genomes_v1* and *genomes_v2* databases *c*) and *d*) contain 155 and 172 species (one genome per species) covering 35 and 20 fish families, respectively. Thus, while the extended *genomes_v2* database contains fewer families, it is more targeted at the diversity present in the Ave River (more species).

To test our ability to distinguish between true and false positives, and between true and false negatives, the *transcriptomes* and both *genomes_v1* and *genomes_v2* databases included species known to occur at the sampling site (whenever available; i.e., included as potential true positives if they are present in the eDNA sample) but also all other species from all fish families known to occur in Portuguese freshwater systems (Collares-Pereira et al., 2021) with available transcriptome and genome sequence data. Additionally, marine fish families commonly present in human diet (e.g., Gadidae, Moronidae) were included in the databases since their eDNA may be present in the water samples. Finally, several additional species not expected to have any match were added as negative control. In case no genome or transcriptome sequences were available for a given target fish family (as in families Cobitidae, Atherinidae, and Loricariidae), members of the next higher taxonomic group were included. In contrast, given the large number of genomes available for Cichlidae (>500), only species from the subfamily Cichlasomatinae, with one representative introduced to Portugal (Chameleon cichlid, *Australoheros facetus*), were included to save computational resources.

Possible contaminants in databases *b*) to *d*) were detected with Kraken2 (Wood et al., 2019) by comparing the sequences with common contaminants plus some curated fish genomes. Putative contaminants in the genomes were surveyed using the following kraken databases: bacteria, archaea, viruses, protozoa, fungi, human, plasmid, and UniVec. Genome sequences from 18 fish species were included to attract specific matches (Table S2). Since non-assigned scaffolds can contain contaminants as well, only scaffolds categorized as chromosomes from these 18 species were included. Genome sequences matching any non-fish taxa were excluded from the assemblies using the extract_kraken_reads.py script from KrakenTools (Lu et al., 2022). The resulting genome sequences were compiled in a final BLAST database where low complexity regions were masked with DustMasker (Morgulis et al., 2006).

### 2.7 Taxonomic assignment of metagenomics reads

Taxonomic assignment of metagenomics sequence reads was performed by megablast (Korf et al., 2003) searches against the four reference databases, using both the merged and unmerged reads >50 bp. For the latter, the read 1 and read 2 files were blasted separately and combined afterwards. BLAST searches only considered hits with a minimum identity value of 90%, query coverage of 90% and E-value of 1^-10^. These were saved in tabular format containing the following information: query id, subject id, taxonomic id, percentage of identity, query length, alignment length, E-value, and bit score.

Outputs from megablast were processed in four consecutive steps using custom scripts available at *github* (https://github.com/mcurto/eDNA_Metagenomics) to obtain a final taxon list from the samples. These steps consisted on 1^st^. combining information of hits from unmerged reads, 2^nd^ selecting the best matches for each read, 3^rd^ extracting the lineage information for each match; and 4^th^ summarizing the lineage information up to the most recent common ancestor across all matches. Details can be found in supplementary methods.

In addition to the classification for each database, we combined the results from the four databases by finding the best hit per read across them. In cases of multiple best hits, the most recent common ancestor was taken as the final taxon. The SeqIDist approach was then implemented to exclude false positives by considering sequence identity distributions with a mode below 99.5% as false positives.

Computational analyses were performed on the Saga Cluster owned by the UiO and the Norwegian metacenter for High Performance Computing (NOTUR) and operated by the UiO Department for Research Computing (https://www.hpc.uio.no).

### 2.8 Filtering metagenomic matches to minimize database contaminants

Despite filtering out contaminants in genome assemblies, BLAST matches of the “cleaned read data” still showed suboptimal hits solely to non-fish taxa, namely bacteria. Therefore, as an additional quality control criterion, only reads matching fish sequences in the *nucleotide* database *a*) were considered for downstream analyses. This decision is based on the expectation that, given the large number of genomic resources for fish species included in the reference database, any read matching a fish locus should match an orthologous region from other fish taxa with lower sequence identity.

In a preliminary experiment, we observed that highly conserved regions, such as some ribosomal loci and low-complexity regions, can generate false positive species detections, especially when reference databases are incomplete. These conclusions were drawn from mapping the metagenomics reads to two fish genomes: *Anguilla anguilla* (GCF_013347855.1) that is present in the study river system, and *Oreochromis niloticus* (GCF_001858045.2) that is absent. We assume that genomic regions with higher-than-average read mapping depth are hotspots for unspecific matches, which was the case for ribosomal loci and low-complexity regions (data not shown). Based on these observations, low-complexity regions were masked in the reference databases and reads matching ribosomal loci in the *nt* database were excluded from analyses.

### 2.9 Reciprocal and recursive BLAST comparisons

For three genera (*Gobio*, *Luciobarbus*, and *Squalius*) we found matches in the metagenomics data for closely related species. In principle, differences in assembly quality among very closely related species may contribute to false positives. Fragmented and incomplete assemblies of species that are truly present may fail to attract some of their conspecific matches, which would in turn align with high identity to a nonspecific better-quality interspecific assembly. If that is the case, we expect that reads matching the higher quality assembly (i.e., an assembled region missing in the assembly of a close relative) will not match the lower quality one. To test this hypothesis, we conducted reciprocal BLAST of the eDNA reads detected between genome assemblies of species pairs where this might be happening: *G. gobio* (higher quality) versus *G. lozanoi* (lower quality); *L. bocagei* versus *L. comizo* (both low quality); *S. cephalus* (higher quality) versus *S. squalus* (lower quality); *S. cephalus* (higher quality) versus *S. carolitertii* (lower quality).

### 2.10 DNA metabarcoding with 12S mitochondrial gene

We evaluated the sensitivity of fish species detection using eDNA metagenomics by comparing its performance versus the current “gold standard” produced by eDNA metabarcoding, using the MiFish-U primers from (Miya et al., 2015). Amplicon library preparation was done with the dual index strategy similar to (Kozich et al., 2013) with a protocol adapted to the primers used (Supplementary methods).

The obtained reads were used to define amplicon sequence variants (ASVs) and taxonomic assignment was done via BLAST (Supplementary methods). Errors originating from index swapping were minimized by applying a correction factor as described by (Hambäck et al., 2021). We also used the Bayesian approach described by these authors to define variable thresholds of minimum number of reads. The sequencing run included eDNA samples from marine ecosystems amplified using the same marker. This experiment integrated an Illumina run including eDNA samples from marine environments using the same marker. To minimize the false detection of marine taxa due to index swapping, organisms detected in the Ave River samples with higher read numbers in the marine samples were excluded remaining only matches to species that spend part of their life cycle in freshwater as likely true positives.

### 2.11 Read length as a lens into residency time and source of eDNA

The DNA found in environmental samples is a mixture of molecules from different organisms and sources, with variable lengths depending on the time and agent(s) of decay (Jo & Yamanaka, 2022). Upon release of intact DNA molecules into the environment, DNA fragmentation is expected to increase over time, whereby recently shed eDNA will consist of longer molecules (and read lengths) compared to older eDNA (Jo et al., 2017). Beyond residency time, the level of eDNA decay (and read length) may vary depending on the source organisms: living microorganisms will be present in any environmental sample and will contribute intact intracellular DNA molecules to the eDNA sample (Pathan et al., 2021). Their eDNA will exhibit longer read lengths than extracellular eDNA originated from lysed cells. Finally, environmental samples may include DNA of organisms from habitats different from the one sampled. Freshwater systems receive input material from the whole drainage area (i.e., of terrestrial origin) as well as from airborne material (Câmara et al., 2021; Deiner et al., 2016). Thus, eDNA originating from habitats other than the one sampled will undergo decay during transport (Jo et al., 2017). Decay is faster in larger molecules than in shorter ones, meaning that size heterogeneity may be greater in eDNA fragments released more recently into the environment than in those residing for longer periods (Jo et al., 2017). Accordingly, read length standard deviation (SD) of matches to microbes and freshwater organisms should be higher than that of matches to macroscopic and marine taxa. These expectations were tested by comparing read length across taxa and habitats of origin.

All taxa detected with identity distributions congruent with being true positives were classified according to size as either microorganisms (all unicellular organisms, or planktonic multicellular organisms) or macroorganisms (all other), and according to habitat as freshwater, terrestrial, or marine. Some organisms may be found in multiple habitats and were not considered in the analysis. Very few organisms from brackish and airborne environments were detected (Table S2), thus they were excluded to prevent low sample size-biases in the analyses. All these classifications were based on literature search using known databases (e.g., https://www.algaebase.org/, <u>WoRMS - World Register of Marine Species</u>, ncbi, gbif) and google scholar searches using organism name as query. For (microbial) taxa represented by specific strains, the information was taken at the strain-level. Taxa missing information in more than one variable were not included in the analysis. Read length information taken from matches to all databases was retrieved for statistical analysis. Models for read length are Generalized Linear Mixed Models (GLMMs) fitted with Gaussian distributions and with species identity included as random intercept factor. Models for read standard deviation are Generalized Linear Models (GLMs) fitted with Gaussian distributions. Models were fitted in R (Team, 2021) using the function *lmer* of lme4 package (Bates et al., 2014) for GLMMs and the function *glm* of stats package (Team, 2021) for GLMs.

## 3 Results

### 3.1 Accuracy of multilocus SEQIDIST taxon identification at species level

As a proof-of-principle, we used publicly available shotgun whole-genome sequence data (sgWGS) from different fish species to show that sequence identity distributions of blast hits against assemblies of conspecifics display a highly left-skewed distribution peaking at 100% sequence identity. In turn, as genetic distance between species increases, BLAST hits against heterospecific genomes peak at <99.5% sequence identity. These expectations were tested through 139 comparisons of short read data from genomic DNA against reference genomes (Table S1). From an initial 1000 genomic DNA reads used per sample, between 85 to 966 reads matched a certain reference genome with at least 90% identity. All intraspecific comparisons resulted in distributions with a mode at 100% identity, the true positives, as most reads of a species did not differ from its reference genome (Figure 1, Figure S1). Most interspecific comparisons, however, resulted in distributions with a mode below 100% (Figure 1, Figure S1). The exceptions being: *Luciobarbus bocagei* vs. *L. comizo*, *Micropterus salmoides* vs *M. nigricans*, *Salmo trutta* vs. both *S. ischchan* and *S. caspius*, *Squalius carolitertii* vs. *S. pyrenaicus*, and *S. squalus* vs. *S. cephalus* (Figure S1).

**Figure 1.**
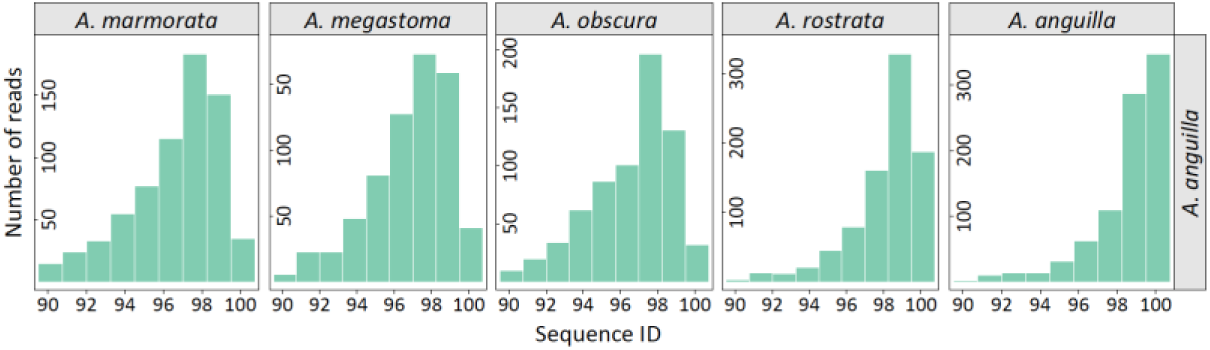
Intra- and interspecific identity distributions of 1000 shotgun whole-genome sequences. Short-reads of *A. anguilla* (SRR12854530) are compared to the genomes of five eel species. The intraspecific comparison (right) shows a distribution with mode at 100% sequence identity, while interspecific comparisons display modes <99.5% and are less left-skewed. For additional comparisons see Figure S1.

### 3.2 Whole community biodiversity assessment

Of the 31,519,078 shotgun sequencing reads produced, 84.5% (26,627,319) passed the quality control and 82.1% (21,855,943) were successfully merged (Table S3). Nevertheless, only 4% (1,167,349) and 0.8% (210,547) matched the *nt* database at the >90% and >99% identity thresholds, respectively. The SEQIDIST approach recovered 1933 taxa, from 73,579 reads with identity above 99%, across all domains of life (Figure 2), encompassing 66 phyla, 427 families, and 1413 species of micro- and macroscopic organisms (Table S2). Most of the reads are assigned to bacteria (77.7%), with 21.4% assigned to eukaryotes, and 0.63% and 0.24% to archaea and viruses, respectively (Figure 2). Within Eukarya, we detect 12 animal (metazoan) phyla (25 classes, 54 families, 65 genera), 2 green plant phyla (8 classes, 58 families, 81 genera), 5 fungi phyla (14 classes, 26 families, 26 genera), and 11 protist phyla (24 classes, 76 families, 108 genera). Microalgae (e.g., Bacillariophyta and Cryptophyceae) were the most abundant eukaryotic taxa detected. We detect not only freshwater organisms but also taxa from nearby marine (20 phyla, 31 classes, 49 orders, 56 families, and 69 genera), brackish (5 phyla, 8 classes, 10 orders, 10 families, and 9 genera) and terrestrial environments (30 phyla, 57 classes, 118 orders, 182 families, and 316 genera) (Table S2).

**Figure 2.**
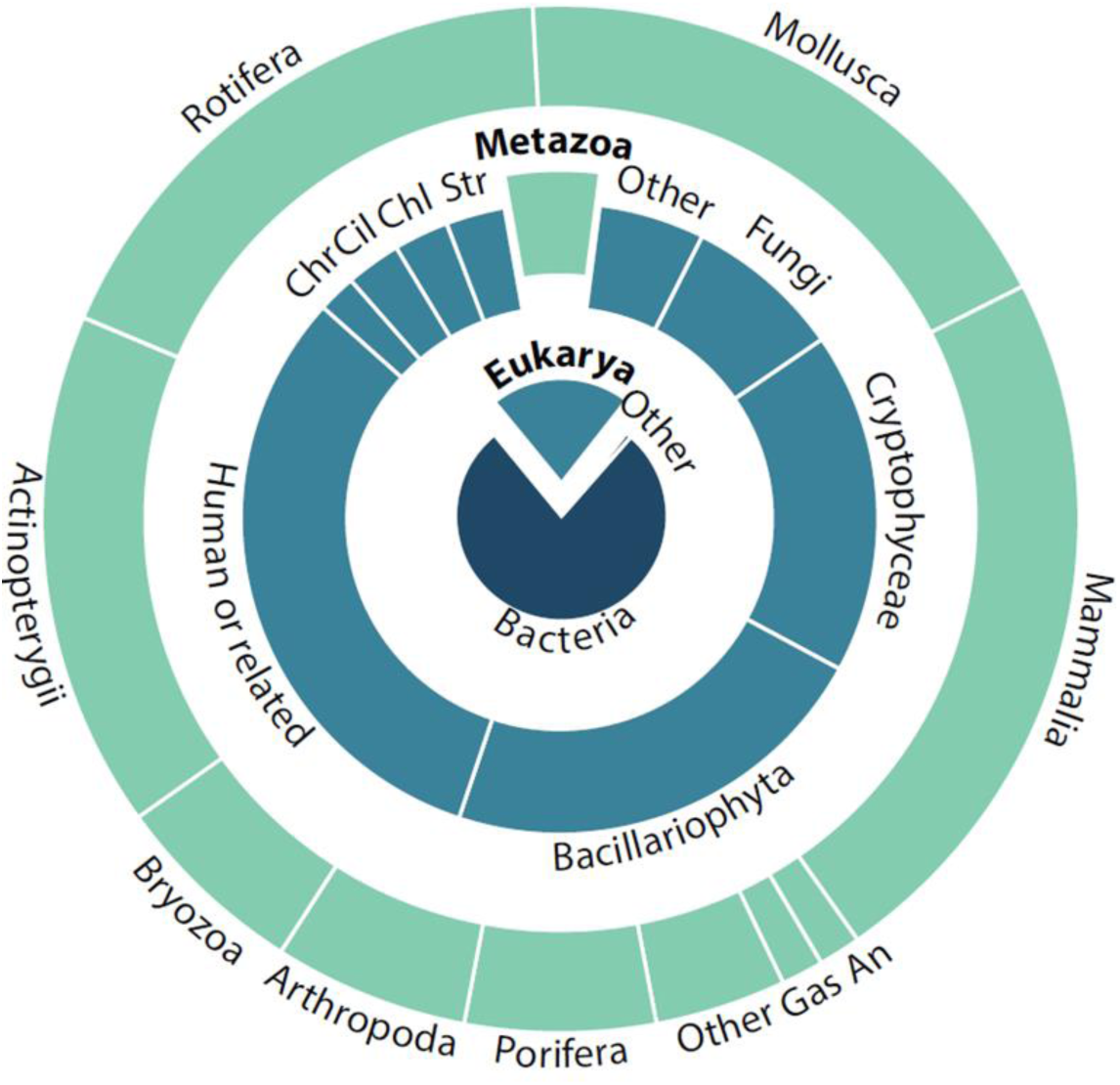
Proportion of metagenomic reads assigned to taxa across the tree of life using the GenBank nucleotide (*nt*) database. Human or related (other primates) are plotted independently from the remaining metazoans to facilitate the visualization of the relative abundances of animal taxa. Some taxon names are abbreviated: Str–Streptophyta, Chl–Chlorophyta, Cil–Ciliophora, Chr–Chrysophyceae, Gas–Gastrotricha, An–Annelida.

### 3.3 Fish diversity and database comparison

Considering the top blast hits of eDNA metagenomics reads against the four databases, 3,346 and 2,342 reads match fish taxa with >90% and >99% identity, respectively. The *genomes_v1* and *genomes_v2* databases provide the highest fraction of the blast hits matching fish taxa, although the extended *genomes_v2* database clearly outperforms the others (96.6%, versus 28.6% with genomes_v1). Indeed, most of the fish reads (67.4%) match only the extended *genomes_v2* database (Figure S2). The *transcriptomes* (9.5%) and *nt* (6.1 %) databases have fewer than 10% of the fish reads producing blast hits above the identity cutoffs.

The combination of all four databases resulted in matches to 151 fish taxa with identity above 90% and 48 above 99%. However, only 21 of these have multilocus identity distributions consistent with true positive detections using SEQIDIST (Figure 3; Table S4). These included three marine species common in local human diet (*Dicentrarchus labrax*, *Gadus morhua* and *Sardina pilchardus*), whose eDNA likely reached the system through wastewater. Thus, while their detection is true, their presence in the riverine system is not.

**Figure 3.**
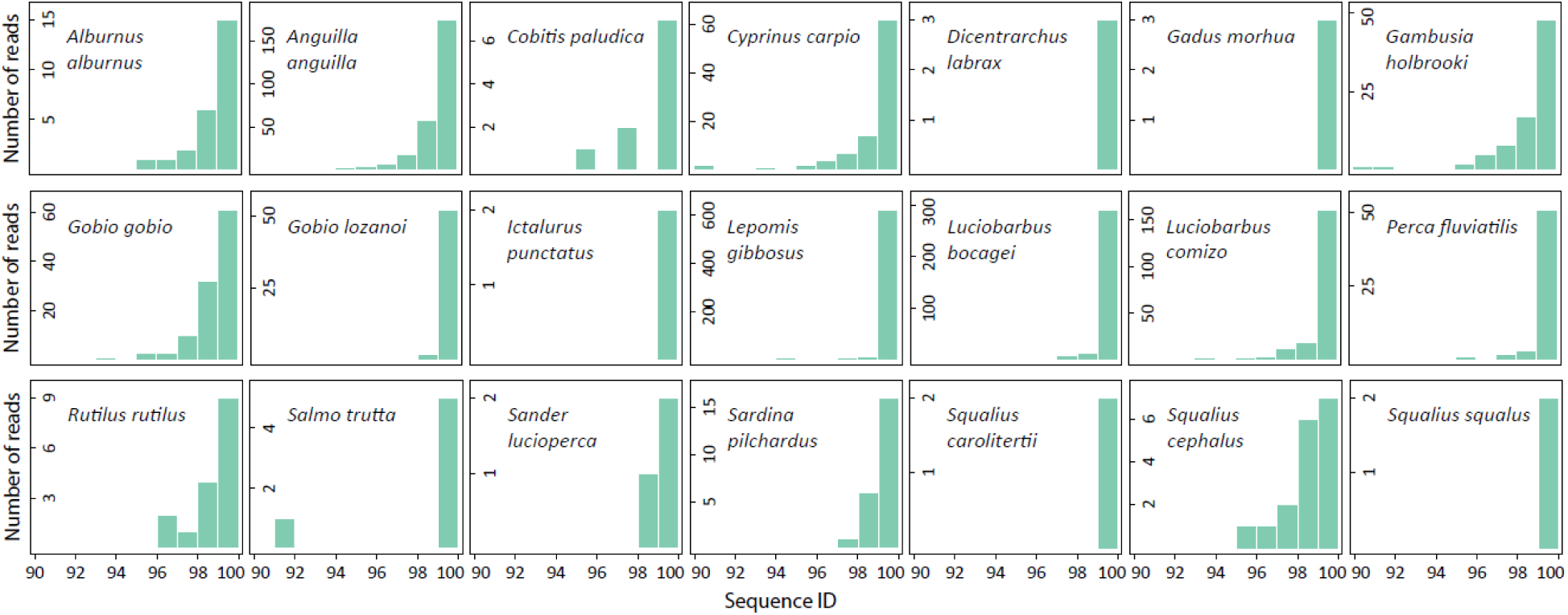
Sequence identity distributions of metagenomics reads matching fish taxa of the Ave River. Only left-skewed distributions with mode at 100% are shown (i.e., congruent with true positives), using the combined result of the four reference databases. Three are marine species common in human diet whose eDNA likely reached the system through wastewater (*Dicentrarchus labrax*, *Gadus morhua* and *Sardina pilchardus*), while *Luciobarbus comizo* and *Squalius cephalus* are likely false positives due to a combination of ILS and introgression.

Three genera (*Luciobarbus*, *Gobio,* and *Squalius*) showed distributions with modes at 100% for two congeners. We conducted reciprocal and recursive blast searches to exclude the possibility of false positives caused by differences in assembly quality between very closely related species. All reciprocal blast distributions had modes at lower identity values, typical of interspecific comparisons (Figure S3). Once we run recursive blasts, in which the reads of the reciprocal blast are remapped to the first genome, the results show similar identity distributions typical of intraspecific comparisons (Figure S4). Altogether, a hypothetical artifact stemming from differences in assembly quality can be excluded, thus ruling-out the possibility that assembly quality is driving these detections.

The intraspecific comparisons with identity distribution modes at 100% included species that have been previously detected for the Ave River using independent methods (*Luciobarbus bocagei*, *Gobio lozanoi*, *Squalius carolitertii, S. squalus*), and taxa that would constitute new records (*Lucobarbus comizo*, *Squalius cephalus and Gobio gobio*). For the proof-of-principle test using sgWGS, intraspecific comparisons with identity distribution modes at 100% were already found for *L. bocagei* vs. *L. comizo*, and *S. squalus* vs. *S. cephalus*. These are likely a consequence of introgression or incomplete lineage sorting (ILS) indicating that one of these species per pair is likely a false positive (Note S1). This was considered to be the case of *Luciobarbus comizo* and *Squalius cephalus* since they are not historically present in the Ave River.

After excluding *L. comizo* and *S. cephalus*, as well as the three marine species, a final list of 16 species detected was obtained with SEQIDIST metagenomics approach for the Ave River samples. All of these but one (*Ictalurus punctatus*) was detected by the *genomes_v2* database (Figure S2, Table S4) outperforming the *genomes_v1* that detected only 10 species. Both *genomes* databases outperformed the *nt* and *transcriptome* databases that retrieved only five and four taxa, respectively.

The addition of new genomes in the *genomes_v2* database reduced the number of matches at lower identity in distributions of related species. This was evident in *Cyprinus carpio*, that had a multimodal distribution for the *nt* and *genomes_v1* databases and a unimodal distribution peaking at 100% for the *genomes_v2* database (Figure S5). Another example was the case of *Micropterus nigricans* (Centrarchidae) that, at the *genomes_v1* database, showed an identity distribution congruent with a false positive (ID mode = 93%) with a high number of matches (Figure S5). By including the *L. gibbosus* (Centrarchidae) genome in our extended *genomes_v2* database, the number of reads matching the *M. nigricans* genome (even with a more complete assembly) is greatly reduced while those matching *L. gibbosus* show a highly skewed identity distribution centered at 100% identity.

### 3.4 Fish eDNA metabarcoding results

The MiFish-U marker was used as the “gold standard” of eDNA analysis to assess the relative performance of metagenomics. Around 1.4M reads were obtained, of which 494,926 passed all quality control, denoise and chimera filtering steps, resulting in 198 ASVs. 103 ASVs are assigned to fishes (464,495 reads) from which 95 are kept after correcting for index swapping. These ASVs match 40 fish taxa, of which 24 are marine taxa that exhibited the highest read numbers in the marine samples sequenced in the same run as the Ave River samples. These matches likely result from index swapping or DNA may originate from wastewater (Table S5). Therefore, we only consider matches to species that spend part of their life cycle in freshwater as likely true positives, resulting in 14 fish species detected in the Ave River samples (Table S4 and S5).

### 3.5 eDNA residency time inferred from read length distributions

To test the hypothesis that read length could serve as a predictor of eDNA source, with longer residency times and greater transport distances in the environment expected to result in more decayed eDNA, we categorized all taxa detected in the matches according to their size and native habitat, resulting in: microorganisms-1378 taxa and 44242 reads; and macroorganisms-205 taxa and 6027 reads; freshwater - 570 taxa and 30,347 reads; marine-100 taxa and 2188 reads; terrestrial-682 taxa and 12,852 reads; brackish-10 taxa and 423 reads; and airborne: 1 taxon and 1 read (see Table S2 for details). Because of their low numbers, brackish and airborne taxa were not considered in further analyses. Habitat information was either not found or reported as multiple habitats for 570 taxa (27,768 reads), while size information was not found or was ambiguous for 350 taxa (73,579 reads). The final datasets of read lengths used in the following analyses can be found as supplementary data (Supplementary Data 6 to 7).Using Generalized Linear Mixed Models (GLMMs) for read length and Generalized Linear Models (GLMs) for read SD, we find that reads assigned to microorganisms are significantly longer and have more heterogeneous lengths (mean = 253.7, SD = 118.6) than those of macroorganisms (mean = 190.7, SD = 105.0), while reads assigned to freshwater organisms are longer and more heterogeneous in length (mean = 264.5, SD = 119.0) than those assigned to marine (mean = 258.1, SD = 116.1) and terrestrial (mean = 210.5, SD = 115.9) organisms (Figure 4; Table S6).

**Figure 4.**
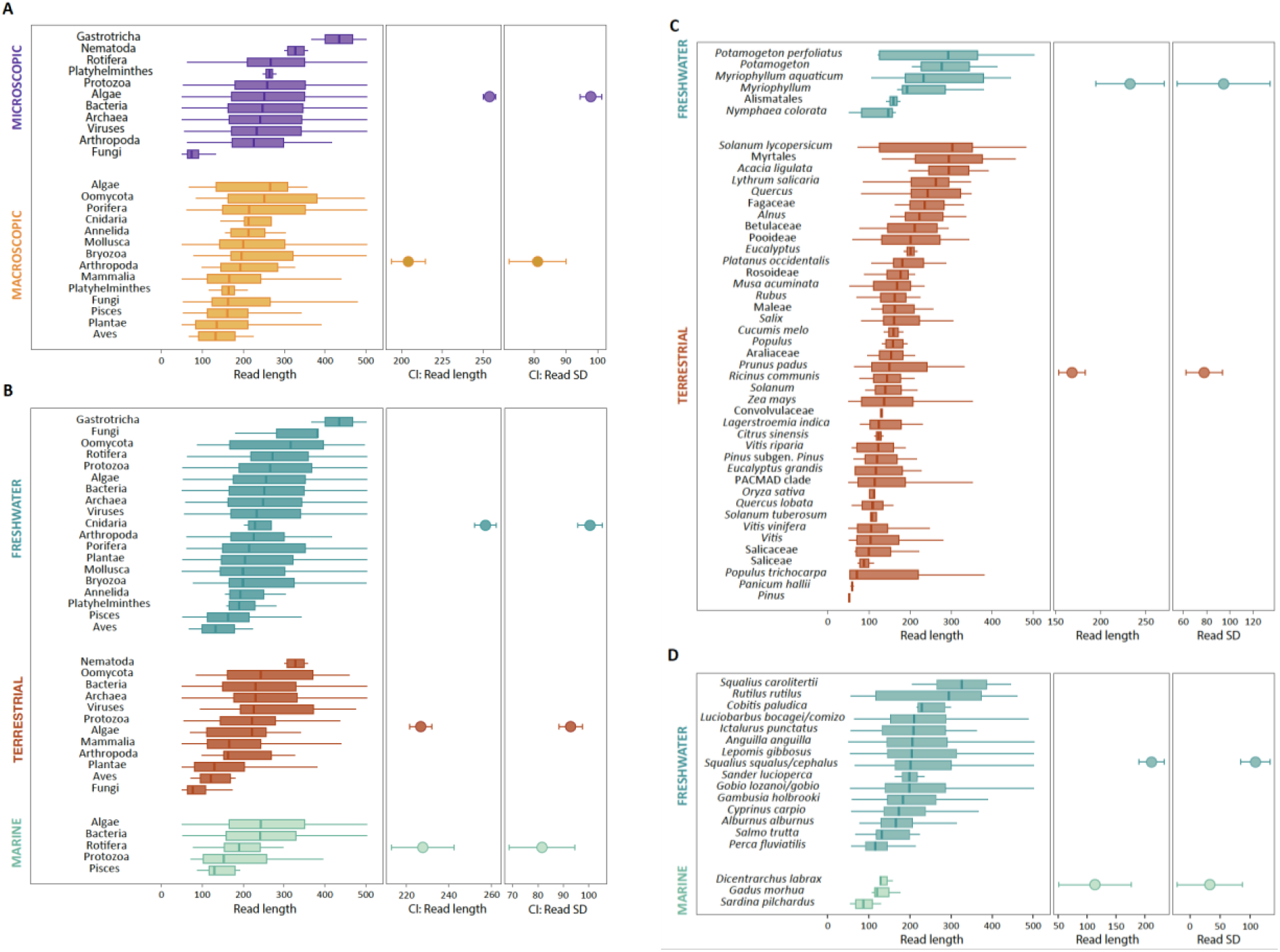
Read length and standard deviation across groups and species. Comparisons between (A) macroscopic vs microscopic organisms; and different habitats across (B) all organisms; (C) terrestrial plants, and (D) fishes. Read length per taxa included in each category is summarized in the left panel by boxplots with median (middle bar), inter-quartile range (box) and extremes up to 1.5 times the inter-quartile range (whiskers). Middle and right panels show 95% confidence intervals based on a Generalized Linear Model (GLM) and Generalized Linear Mixed Model (GLMM), respectively. Model statistics are presented in Table S6.

A similar effect of native habitat is found for plant taxa (Figure 4; Table S6). Reads assigned to freshwater plants (12 taxa and 41 reads) are longer and have more heterogeneous lengths (mean = 241.6, SD = 124.1) than those matching terrestrial plants (82 taxa and 411 reads; mean = 153.8, SD = 92.4). Among terrestrial plants, taxa used in agriculture tend to show lower read length (mean = 140.8, SD = 86.9) compared to taxa naturally found around and along the riverbank (mean = 173.4, SD = 97.6), although the differences are not statistically significant (Figure 4; Table S6). While eDNA from wild terrestrial plants is integrated into the aquatic system through remains of individuals growing nearby, eDNA of agricultural taxa is more likely to have been transported through the sewage system in addition to going through digestive systems, and thus residing in the environment for a longer time.

Considering the specific case-study of fish, reads assigned to freshwater taxa show significantly longer and more heterogeneous lengths (14 taxa and 1577 reads; mean = 223.8, SD = 107.9) compared to reads assigned to marine taxa (3 taxa and 24 reads; mean = 107.5, SD = 43.9). Within freshwater taxa, *Salmo trutta* (mean = 148.0, SD = 62.8) and *Perca fluviatilis* (mean = 116.9, SD = 37.0) also have shorter read lengths, which may be due to increased eDNA degradation associated with transport from upstream populations and thus reflect longer residency times (Figure 4).

## 4 Discussion

### 4.1 Excluding false positives with SEQIDIST

SEQIDIST solves the high prevalence of false positives of ‘traditional’ metagenomics when classifying at the species level (Simon et al., 2019). Taxon detection and discovery using individual sequence matches ultimately leads to unacceptably high rates of false positives, while simultaneously overseeing false negatives (e.g., fishes in (Manu & Umapathy, 2023)). Alternatively, a ‘pseudo-taxonomic approach’ could be implemented, where read classification is performed as putative molecular operational taxonomic units (Cowart et al., 2018), or as classifications to the closest higher taxon, making taxonomic assignments at higher taxonomic levels (e.g., genus; Manu and Umapathy, 2023). However, that is done at the cost of information loss and increased taxonomic composition uncertainty (increasing both false-negative and false-positive assignments). Additionally, matches to erroneous taxa can occur even at high taxonomic levels (i.e., families, orders) with deceiving statistical confidence in conserved genomic regions of high sequence identity, thus not avoiding false positives despite trading-off taxonomic certainty.

The SEQIDIST framework takes all blast hits to each individual taxon to make a species identity profile. By using sgWGS data from different fish species we demonstrate that profiles resulting from conspecifics peak at 100% sequence identity, contrasting with heterospecific comparisons that peak at lower values. Such property of identity distributions makes false positives become easily discernible. However, some interspecific comparisons had a distribution profile with a mode at 100%. All these correspond to very closely related species, hybrid species, and species for which natural hybridization has been reported to be common (e.g. Gante et al., 2015; Gouskov & Vorburger, 2016; Segherloo et al., 2021)). Thus, if the individual used for building a reference genome has been introgressed, its genome will show an enrichment of high identity reads between the hybridizing species. Additionally, incomplete lineage sorting contributes to the retention of ancestral variants, resulting in patterns of shared variation across closely related taxa that do not match the species tree (Pollard et al., 2006). Therefore, the high sequence identity found between some closely related species is likely a consequence of shared genomic variation caused either by ILS or introgressive hybridization. In such cases, there is the potential for false positives, but this can be at least partially controlled for.

Nevertheless, possible misdetections would happen mostly in the scenario of species discovery, where true species presence could be difficult to disentangle from introgression and ILS, especially in systems with many closely related species. In the case of species monitoring in better known systems using eDNA metagenomics, determination of species identity can take into account other sources of information, such as a pre-existing knowledge of species diversity at the site based on historical sampling data, and knowledge of the phylogenetic relationships between the putative target species co-occurring in the system, or with previous evidence of introgression (e.g., eliminating off-target species with closely-related congeners in the system from databases, or monitoring detection proportions). In practice, this particular effect resembles the impact of low marker resolution, ILS and introgression also observed in (e)DNA (meta)barcoding. Thus, proper vouchering in natural history collections of specimens used to produce the reference nuclear genomes and mitogenomes (Buckner et al., 2021) is a pre-requisite for accurate species identification using eDNA, and to rule-out possible impacts of introgressive hybridization or other marker limitations on molecular information content.

### 4.2 Describing diversity across the tree of life

SEQIDIST accurately describes biological communities across the tree of life. We identified 1933 taxa from all domains, which is at the same order of magnitude of recent eDNA metagenomics studies from more diverse tropical brackish lagoons in India and the Southern Ocean (Cowart et al., 2018; Manu & Umapathy, 2021, 2023), despite our use of more stringent filtering (smaller E-value and higher sequence identity). Such relatively high number of taxa was primarily due to the ability of SEQIDIST to confidently assign matches at species levels. Nevertheless 96% of reads were not assigned at all, highlighting the amount of biodiversity information still unmapped in the generated dataset. Most matches found were to microorganisms, such as bacteria and diatoms that are generally abundant and largely represented by intracellular eDNA that is favourably sequenced (Singer et al., 2020; Stat et al., 2017). In addition, bacterial eDNA tends to be more easily assigned to taxa due to their low complexity and better representation in reference databases (Singer et al., 2020; Stat et al., 2017). These proportions may also reflect the high degree of eutrophication at the sampling site (A. Veríssimo and G. Riccioni, pers. observation), which favours microbial activity.

Much of the diversity detected was from non-freshwater organisms, whose presence can be explained by multiple processes. Terrestrial eDNA can be originated not only from species that contact the river directly but also from biomass that is added to the river through sewage systems (Macher et al., 2021). This assessment is supported by the high abundance of human eDNA and the detection of organisms that are commonly found in wastewater, used for human consumption, or associated with agriculture and farming (Table S1). For instance, the detection of some marine vertebrate taxa (e.g., marine fishes) in freshwater samples is likely attributable to human consumption; however, occasional intrusions of some marine species, such as the European seabass (Almeida et al., 2024), may also occur. Other sources of eDNA reflect the wide spatial scale of biodiversity represented in freshwater systems. For instance, marine microalgae have been detected in freshwater lakes and other terrestrial environments near the ocean due to the aerosolization of sea surface water and inland wind transport (Câmara et al., 2021). In fact, many organisms are part of the so-called bioaerosols and can be transported for long distances (Razjigaeva et al., 2022). In our study, this phenomenon is exemplified by the detection of microalgae like *Thalassiosira pseudonana* (Bacillariophyta) or *Crustomastix sp.* (Chlorophyta), that are marine taxa according to www.algaebase.org. Given the proximity of the sampling site to the Atlantic Ocean (∼5 km in a straight line), it is plausible that the detection of marine taxa originated from marine bioaerosols.

### 4.3 Discovering new diversity with SeqIDist

SEQIDIST ability to exclude false positives enhances confidence in the discovery of new diversity. Of the final set of 16 species, most have been previously reported to the Ave River, but four are novel (Collares-Pereira et al., 2021), constituting new alien species for the basin (*Ictalurus punctatus, Perca fluviatilis*) and even for the Iberian Peninsula (*Gobio gobio*, *Squalius squalus*). Their detection is not surprising, as this drainage has been a hotspot for illegal fish introductions (Ribeiro et al., 2009; Ribeiro & Veríssimo, 2014), highlighting the power of metagenomics in detecting alien species at the invasion front, generating strong hypotheses regarding species distribution and community composition. If only individual matches were considered, as many as 151 fish taxa would be detected, which would represent an unacceptably high false positive rate. Even when considering only stringent BLAST hits with >99% sequence identity and >90% read coverage, the number would decreased to 48, many of which would still be false positives. Given such uncertainty, it would be impossible to confidentiality report these new occurrences to the Ave River.

The focus on fish species detection in the Ave River also unravelled the higher sensitivity of eDNA metagenomics compared to metabarcoding. eDNA metagenomics recovered 16 taxa compared to 14 using metabarcoding. The increased sensitivity of metagenomics likely reflects known limitations of metabarcoding, namely false negatives due to differences in priming efficiency across taxa (Krehenwinkel et al., 2017), PCR failure due to eDNA decay (Jo et al., 2017) since 57% of the metagenomics reads classified as fishes are shorter than the 172 bp-long barcode used, or low abundance of the barcode locus in the eDNA mix since the probability of detecting a specific mitochondrial locus is lower than anonymous nuclear regions (Note S2). Nevertheless, there are four species detected exclusively with the metabarcoding assay: *Achondrostoma oligolepis*, *Lampetra fluviatilis*, *Esox lucius*, and *Alosa* sp. (Table S4 to 5). These constitute potential false negatives in our metagenomics dataset, likely resulting from a combination of low eDNA abundance (as indicated by the lowest read counts in metabarcoding) and poor genome quality in the *genomes_v2* database (with an N50 < 5 Kbp for *A. oligolepis* and *Lampetra* spp.). The detection of *A. oligolepis* and *Alosa* sp. is not surprising, given the possible occurrence of the species in this region (Collares-Pereira et al. 2021). However, the occurrence of *Lampreta* sp. And *E. lucius* constitute new records for this Portuguese drainage.

### 4.4 Using read length as a proxy for decay

We detected a wide range of organisms, ranging from micro- to macroorganisms, as well as detections from nearby marine, brackish, and terrestrial environments, whose eDNA molecules are expected to have different residency times and, consequently, varying degrees of decay. This allowed us to test whether read length distributions from metagenomics can be used to discriminate between local and regional transport scales and their associated residency timescales. We confirmed our expectations that eDNA of microorganisms and of freshwater organisms is on average longer and more heterogeneously fragmented. Conversely, eDNA of macroscopic organisms, and of marine and terrestrial taxa originating outside the freshwater system, is shorter and more homogeneously fragmented. These general trends are clearly illustrated by the two best known taxonomic groups of macroscopic eukaryotes in the Ave River: plants and fishes. Aquatic plants are represented by longer reads with higher per species standard deviation than terrestrial species. Likewise, marine fish species have shorter read length than freshwater ones. Their DNA likely reached the riverine environment through wastewater, as they are commonly included in local human diet. Moreover, read length distributions are congruent with differences in habitat preference: species restricted to upstream stretches are represented by shorter and more homogeneously fragmented eDNA, likely due to longer residency time in the system and higher decay. This is the case of *Salmo trutta*, whose known populations are found circa 50 km upstream of the sampling locality (F. Ribeiro, unpublished data). Discrimination between local and transported eDNA, and between intracellular and extracellular eDNA, are a major added value of metagenomics data for community characterization at spatial and temporal scales. These have been pointed out as major limitations of eDNA metabarcoding as a biodiversity monitoring tool, since diversity can be overestimated by the detection of taxa absent at the sampling locality (Deiner et al., 2017).

### 4.5 Impact of database type and completeness on SEQIDIST

Adding whole genome sequences greatly improved the detectability of fish communities with eDNA metagenomics. These results highlight the benefit of using reference databases with a good taxonomic representation of whole genome sequences. Nevertheless, genome data is still lacking for most species (Lewin et al., 2018, 2022). When using incomplete databases, eDNA sequence data originating from taxa not represented in the reference database would likely match reference genomes of related taxa resulting in a SEQIDIST distribution with a mode below 100% of sequence identity. Thus, sequence identity distributions to a close taxon provide important information on false positives and dark diversity (false negatives) in environmental samples. This was clearly showed by the decrease in the number of matches at low identity to the *Cyprinus carpio* and *Micropterus nigricans* genomes in the *genomes_v2* database compared to the more incomplete *genomes_v1*. These matches at low identity were converted in high identity matches to newly added genomes of related species. Thus, adding genomes to reference databases (i.e., *genomes_v2* database) illuminates dark diversity, maximizes species detection, and refines identification in eDNA samples. These results highlight the importance of gathering genome sequences for all diversity across the tree of life.

Our results also show that these reference genomes do not need to be of high quality. While differences in assembly quality of reference genomes of close relatives could, in principle, contribute to false positives, we determined that assembly quality is unlikely to play any major role in detection accuracy using reciprocal and recursive BLAST searches against assemblies of varying qualities available from congeners.

### 4.6 Wide applicability and archival potential of eDNA metagenomics

Sensitivity and accuracy in species detection using SEQIDIST improve with whole genome reference databases. Thus, the potential of eDNA metagenomics datasets as repositories of biodiversity information, akin to environmental vouchers, will only increase as more genomic resources become available globally (Ebenezer et al., 2022; Lewin et al., 2018, 2022). These resources will allow a more comprehensive description of past (Kelman & Moran, 1996), present and future biodiversity from eDNA metagenomics, offering a much-needed historical perspective of community diversity changes. eDNA metagenomics can revolutionize whole biodiversity monitoring across the tree of life, speed-up discovery of hidden biodiversity, and ultimately help address the taxonomic impediment with integrative approaches (Zamani et al., 2022). Analytical strategies could provide access to hidden biodiversity by implementing approaches that describe diversity independently of reference databases (Cordier et al., 2019) and further help understand the scale of biodiversity loss the planet is currently facing.

## Acknowledgments

This work was funded by the Portuguese Foundation for Science and Technology (FCT) under the project ENVMETAGENOMICS (PTDC/BIA-CBI/31644/2017). Additional funds were received from FCT through the strategic plans of the research centers MARE (UID/04292/2020), cE3c (UID/BIA/00329/2020), and CIBIO/InBIO (UIDP/50027/2020), and through the project LA/P/0069/2020 granted to the Associate Laboratory ARNET. FCT provided individual contracts to FR (CEEC/0482/2020), and to AV (https://doi.org/10.54499/DL57/2016/CP1440/CP1646/CT0001); MC and GR were supported by research contracts under the scope of ENVMETAGENOMICS. MC work was also supported by the Horizon 2020 Programme under the Grant Number 857251. HFG was also supported by KU Leuven Research Fund grant STG/21/044.

## Data Accessibility Statement

Metagenomic and metabarcoding raw sequencing data is available at European Nation Archive SRA repository associated with the bioproject PRJEB49862. Barcode sequences for the 12S marker were deposited under the accession numbers OV522633 to OV522672. Genome sequences produced in this study and corresponding short read data were deposited in GenBank under the bioproject PRJNA902967. We also provide supplementary data including the sequence identity distribution for all matches found with the four databases and the combination of them (Data S1-S5) and read length information used in the comparison between taxa size scales and habitat (Data S6-S7). These files can be downloaded at https://github.com/mcurto/eDNA_Metagenomics.

All scripts and codes used in the bioinformatic analysis are available and described at https://github.com/mcurto/eDNA_Metagenomics.

## Benefit-Sharing Statement

There are no benefits to report.

## Author contributions

MC, AV, CDS, FR, MJA and HFG contributed to the conceptualization and experimental design of the study. Field work was conducted by AV, GR, and FR, while laboratory work by AV and GR. Sequence data was analyzed by MC, while statistical analysis was performed by CDS. HFG contributed throughout the analysis’s steps. MC and HFG wrote the first draft of the manuscript that was revised and edited by the remaining authors. All authors contributed to the article and approved the submitted version

## Conflict of interest statement

The authors declare that they have no conflict of interests.

## Notes

https://github.com/mcurto/eDNA_Metagenomics

### Competing Interest Statement

The authors have declared no competing interest.

