## Supplementary information for "Improving whole biodiversity monitoring and discovery with environmental DNA metagenomics"

**Table of contents:**


| | |
|---|---|
| Supplementary methods | Page 2 |
| Figure S1.1 - Intra- and interspecific identity distributions for *Anguilla* WGS data | Page 5 |
| Figure S1.2 - Intra- and interspecific identity distributions for *Gambusia* WGS data | Page 6 |
| Figure S1.3 - Intra- and interspecific identity distributions for *Gobio* WGS data | Page 7 |
| Figure S1.4 - Intra- and interspecific identity distributions for Centrarchidae WGS data | Page 8 |
| Figure S1.5 - Intra- and interspecific identity distributions for *Luciobarbus* WGS data | Page 9 |
| Figure S1.6 - Intra- and interspecific identity distributions for *Salmo* WGS data | Page 10 |
| Figure S1.7 - Intra- and interspecific identity distributions for *Perca* WGS data | Page 11 |
| Figure S1.8 - Intra- and interspecific identity distributions for *Squalius* WGS data | Page 12 |
| Figure S2 - Comparison between the four reference databases | Page 13 |
| Figure S3 - Reciprocal blasts | Page 14 |
| Figure S4 - Recursive blasts | Page 15 |
| Figure S5 - Key examples of sequence identity distributions of metagenomics data | Page 16 |
| Supplementary Notes | Page 17 |
| Supplementary References | Page 18 |

### Supplementary methods

#### *Sampling and DNA isolation*

DNA extraction was performed using three different approaches: using 1) the filtrate, 2) its supernatant, and 3) the corresponding pellet after centrifugation. This approach aimed to maximize extraction of the diversity of eDNA molecules from the filters. The overall community composition and taxa detected did not differ among approaches, thus all data were combined in the final analyses. The eDNA extractions were performed using the Qiagen DNeasy Blood and Tissue kit (Qiagen, Hilden, Germany) according to the manufacturer's instructions, except for the initial digestion step, which was as follows: a) in-filter digestion (3 filters), with 1 ml of proteinase K (20 mg/ml) added into each filter, followed by vortexing and incubation at 55º C overnight (~12h), and removal of 10 ml of digested eluate from the filter to be used in the following extraction steps; b) centrifugation of 15 ml of filtrate (2 filters) at 7830 rpm for 1 h, and separation of 13 ml of the supernatant and the 1 ml of pellet, which were digested separately by adding 1 ml and 200 µl of proteinase K (20 mg/ml), respectively, followed by vortexing and incubation at 55º C overnight (~12h). Final elutions were done in 100 µl of EB buffer.

#### *Metagenomic read taxonomic assignment.*

Taxonomic assignment of metagenomic reads followed four steps. The first step combined information of hits from unmerged reads. This was done by recalculating the sequence identity and query coverage considering the matches from both reads. In case both reads matched the same subject, their values were averaged out; otherwise, the matches were kept separate, but query coverage was recalculated considering the nucleotide sum of both reads. In the second step, the best match per read was selected by considering, in this order, the matches with lowest E-value and highest sequence identity and query coverage. Only matches with a minimum identity and query coverage of 90% were considered. For reads matching multiple records with the same BLAST parameters, taxonomic information from all matches was kept. In the third step, lineages from all matches were retrieved using the corresponding taxids with TaxonKit (Shen & Ren, 2021). For some matches, the taxid had to be fetched from NCBI using the corresponding accession numbers. In the last step, the final lineage assigned to a read was summarized. The lineage up to the most recent common ancestor was considered in case of multiple matches.

#### *Metabarcoding amplicon sequencing protocol*

Amplicon library preparation for metabarcoding was prepared/conducted? in two PCR rounds. The first round of PCRs was performed in a final volume of 25 µL containing 12.5 µL of Qiagen multiplex master mix (Qiagen, Hilden, Germany), 0.5 µL of each primer (10 nM), 5 µL of DNA isolate (5 ng/µL) and 6.5 µL of autoclaved water. The following temperature profile was used for the PCR: an initial denaturation at 95º C for 15 minutes, a first round of 11 cycles of 20" denaturation at 98º C, 15" annealing with temperatures starting at 65º C with a decrease of 0.5º C per cycle, and extension at 72º C for 15"; a second round of 29 cycles with the same conditions of denaturation and extension but the annealing temperature was set to 60º C; and a final extension during 5' at 72º C. To decrease the effect of stochastic amplification biases, eight PCR replicates per eDNA extraction were produced. PCR products were cleaned with Ampure beads (Beckman Coulter, Brea, California, USA) prior to the second round of PCR, for the addition of dual-indexes and Illumina adaptors. Indexing was done using Kapa HiFi Hot Start kit (Roche, Basel, Switzerland) following the manufacturer's recommendations, and the following temperature profile: 95º C for 3 minutes, followed by 10 cycles of 95º C for 30 sec, 55º C for 30 sec, 72º C for 30 sec, and by a final extension step at 72º C for 5 minutes. Indexed PCR products were checked for size and yield by agarose gel electrophoresis and cleaned with Ampure beads. The cleaned indexed PCR products were quantified with Nanodrop (Thermofisher, Waltham, Massachusetts, USA), normalized at 15 nM and pooled equimolarly. The final pool was checked for size and quantity on the TapeStation and validated by qPCR using the KAPA HiFi HotStart Kit. Sequencing was performed in an Illumina MiSeq Paired end run for v2 500 cycles with an expected coverage of 10K reads per sample. This run included eDNA samples from other projects originating in seawater and using the same MiFish primers. The indexing step of library preparation and sequencing was performed by the Centre for Molecular Analysis (CTM) from CIBIO-InBIO.

#### *Metabarcoding ASV definition and taxonomic assignment*

Amplification primer and Illumina adapter sequences were removed with cutadapt (Martin, 2011). Reads not containing the amplification primers were excluded. The R library Dada2 (Callahan et al., 2016) was used to define amplicon sequence variants (ASVs). Quality control was done by removing reads with Ns, allowing up to two expected errors, and truncating reads to 150 bp and 120 bp for read 1 and 2, respectively. Default values were used for the remaining steps and an ASV table was exported. The corresponding sequences were blasted to the NCBI nucleotide database and to a custom-made database of 12S sequences from all species that are expected in the region (ENA: OV522633 to OV522672). Only matches with a minimum E-value of $1^{-10}$ and identity of 90% were considered. The best match (lower E-value and identity) between the nucleotide and custom databases was considered. Taxonomic assignments of read matches were based on the ID value: >99% for species-level, >97% at genus level, >95% at family level and >93% for higher taxonomic levels.

### Supplementary Figures

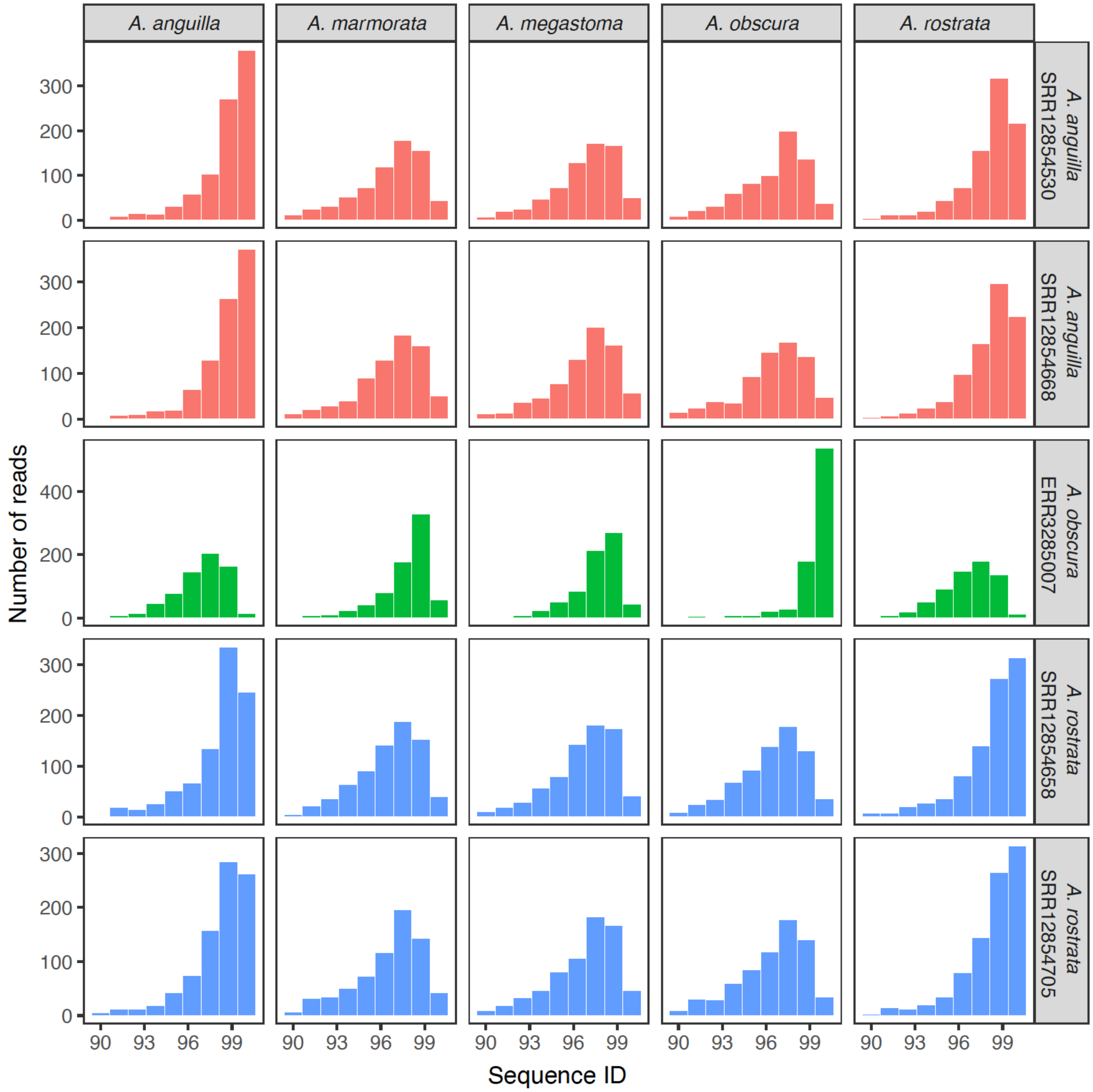


**Figure S1.1.** Intra- and interspecific identity distributions of 1000 shotgun whole-genome sequences of five *Anguilla* species. BLAST searches were performed against three congeneric taxa (accession numbers shown in rightmost column). Intraspecific comparisons generate distributions centered at >99.5% sequence identity and highly left-skewed. Interspecific distributions are centered at lower identities.

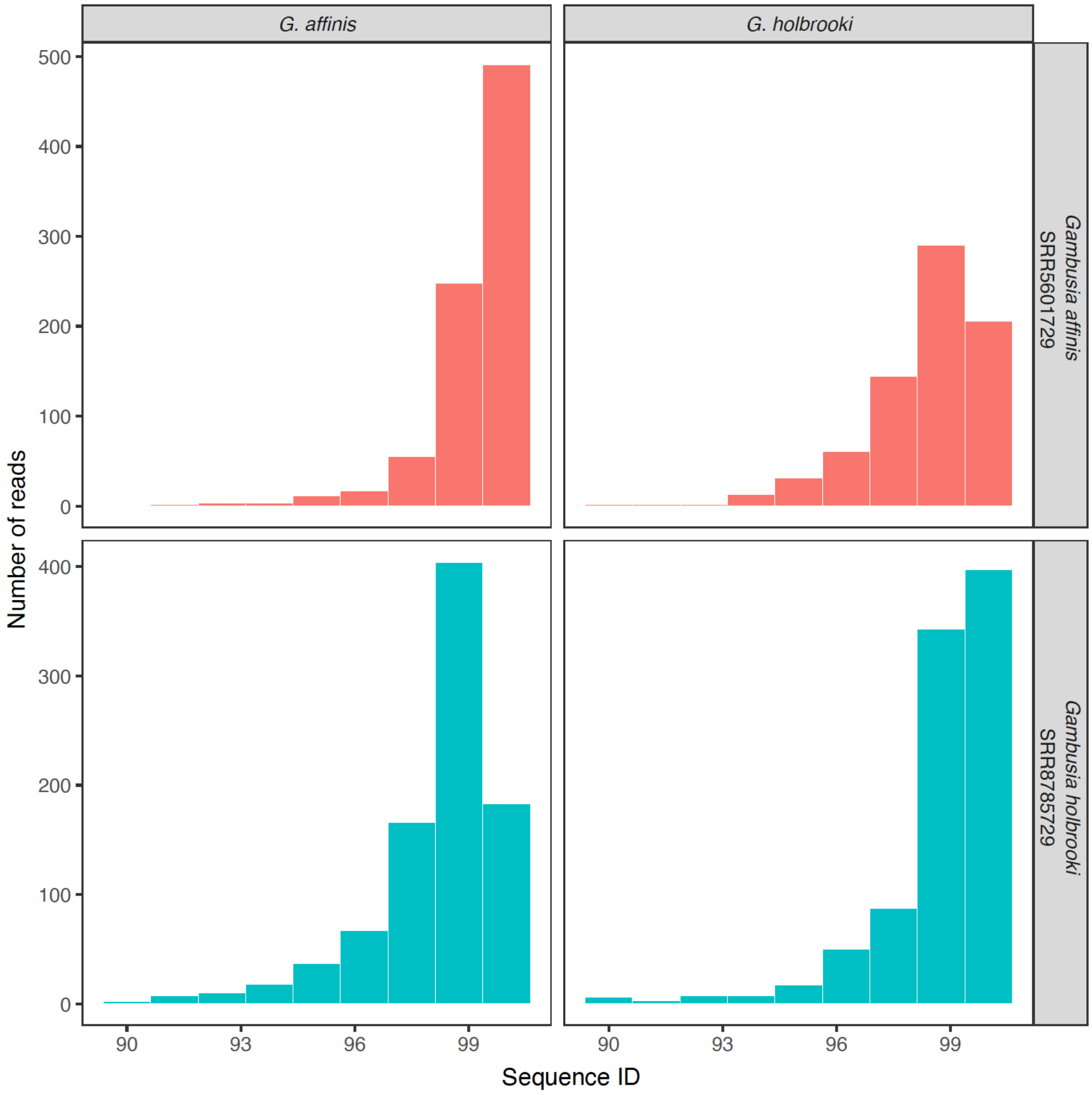


**Figure S1.2.** Intra- and interspecific identity distributions of 1000 shotgun whole-genome sequences of two *Gambusia* species. BLAST searches were performed against each other (accession numbers shown in rightmost column). Intraspecific comparisons generate distributions centered at >99.5% sequence identity and highly left-skewed. Interspecific distributions are centered at lower identities.

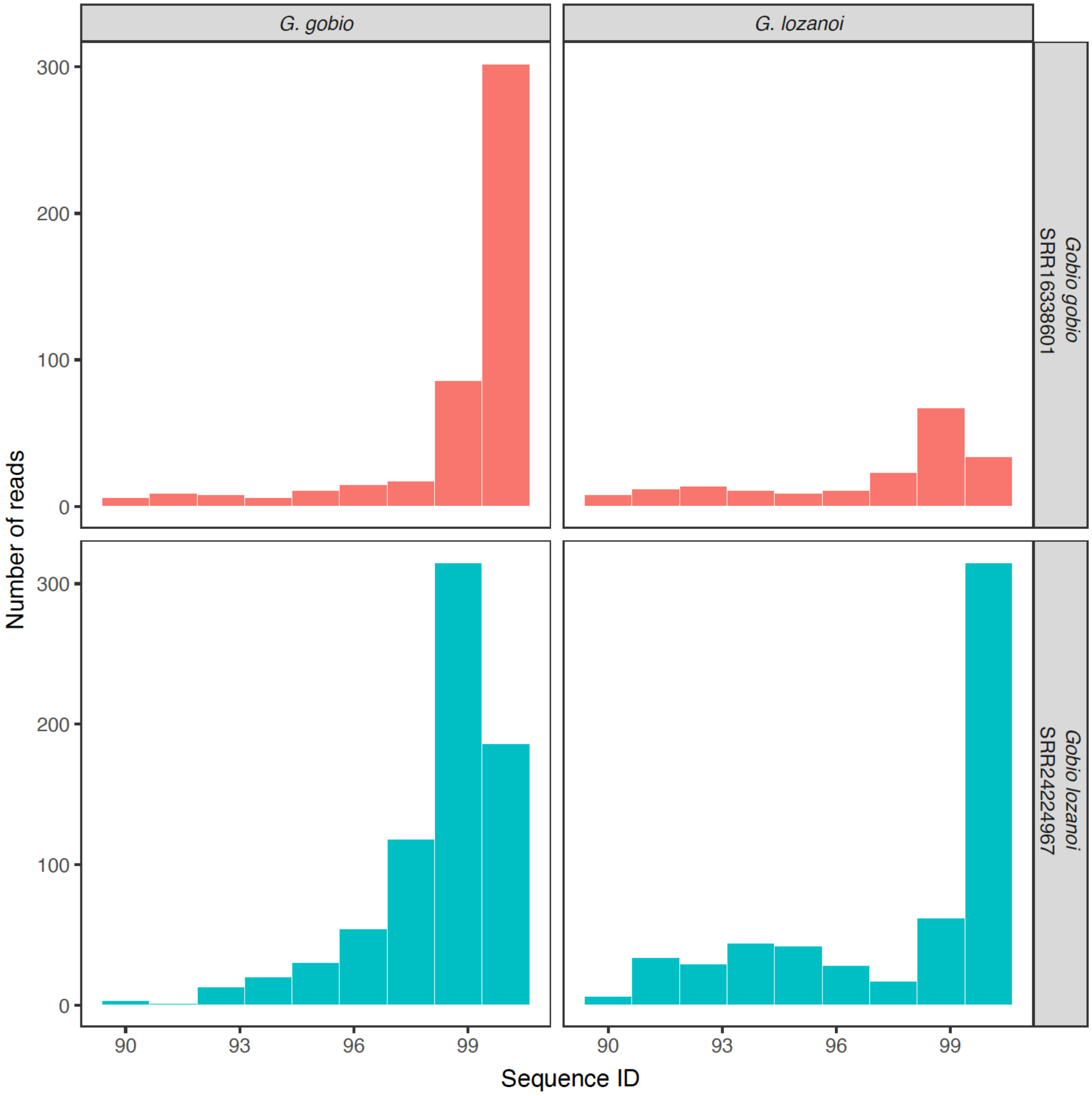


**Figure S1.3.** Intra- and interspecific identity distributions of 1000 shotgun whole-genome sequences of two *Gobio* species. BLAST searches were performed against each other (accession numbers shown in rightmost column). Intraspecific comparisons generate distributions centered at >99.5% sequence identity and highly left-skewed. Interspecific distributions are centered at lower identities.

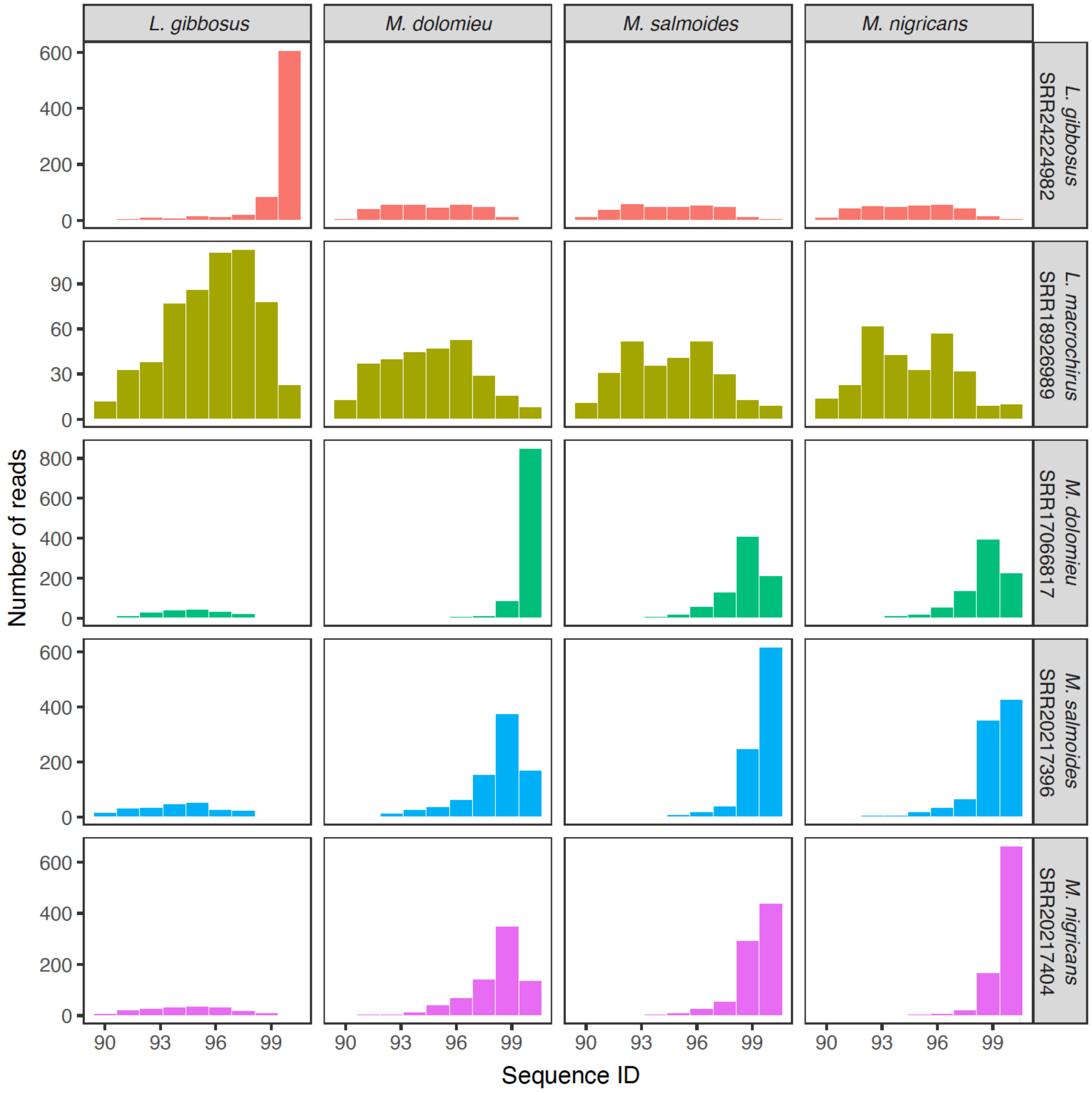


**Figure S1.4.** Intra- and interspecific identity distributions of 1000 shotgun whole-genome sequences of four Centrarchidae species. BLAST searches were performed against each other and to *L. macrochirus* (accession numbers shown in rightmost column). Intraspecific comparisons generate distributions centered at >99.5% sequence identity and highly left-skewed. Interspecific distributions are centered at lower identities with the exception of the pair *M. nigricans/M. salmoides*. *Micropterus salmoides* (formerly *M. floridanus*) has historically been considered a subspecies of *M. nigricans* (formerly *M. salmoides*), with which it displays high ILS and extensive introgression. Nevertheless, note the similar size of the two larger bins, and the lower numbers of matched reads.

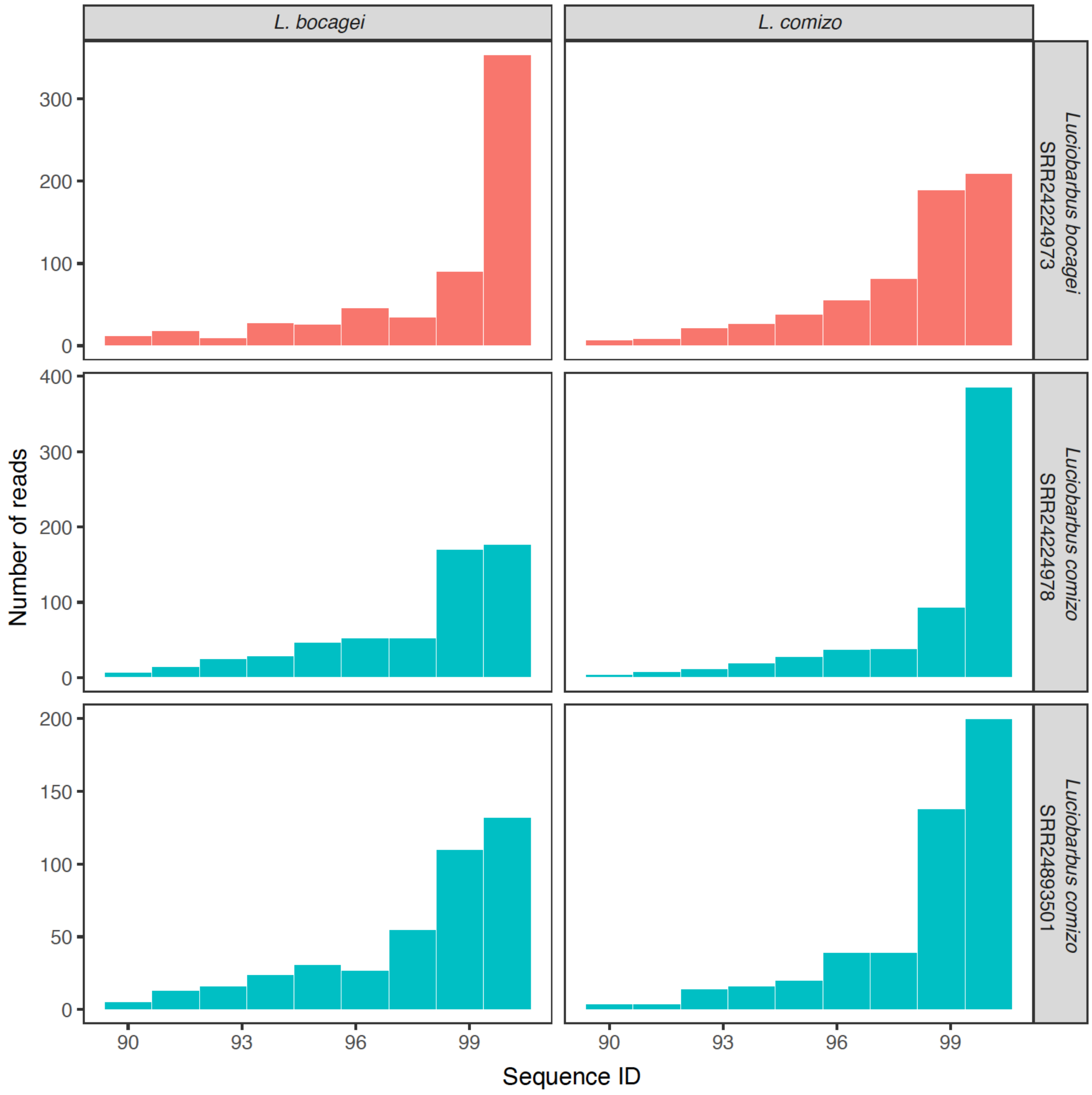


**Figure S1.5.** Intra- and interspecific identity distributions of 1000 shotgun whole-genome sequences of two *Luciobarbus* species. BLAST searches were performed against each other (accession numbers shown in rightmost column). Intraspecific comparisons generate distributions more clearly centered at >99.5% sequence identity and highly left-skewed. Interspecific distributions still have >99.5% modes, but the size of the two larger bins is similar, with lower numbers of matched reads. These species are recently diverged and frequently introgress into each other.

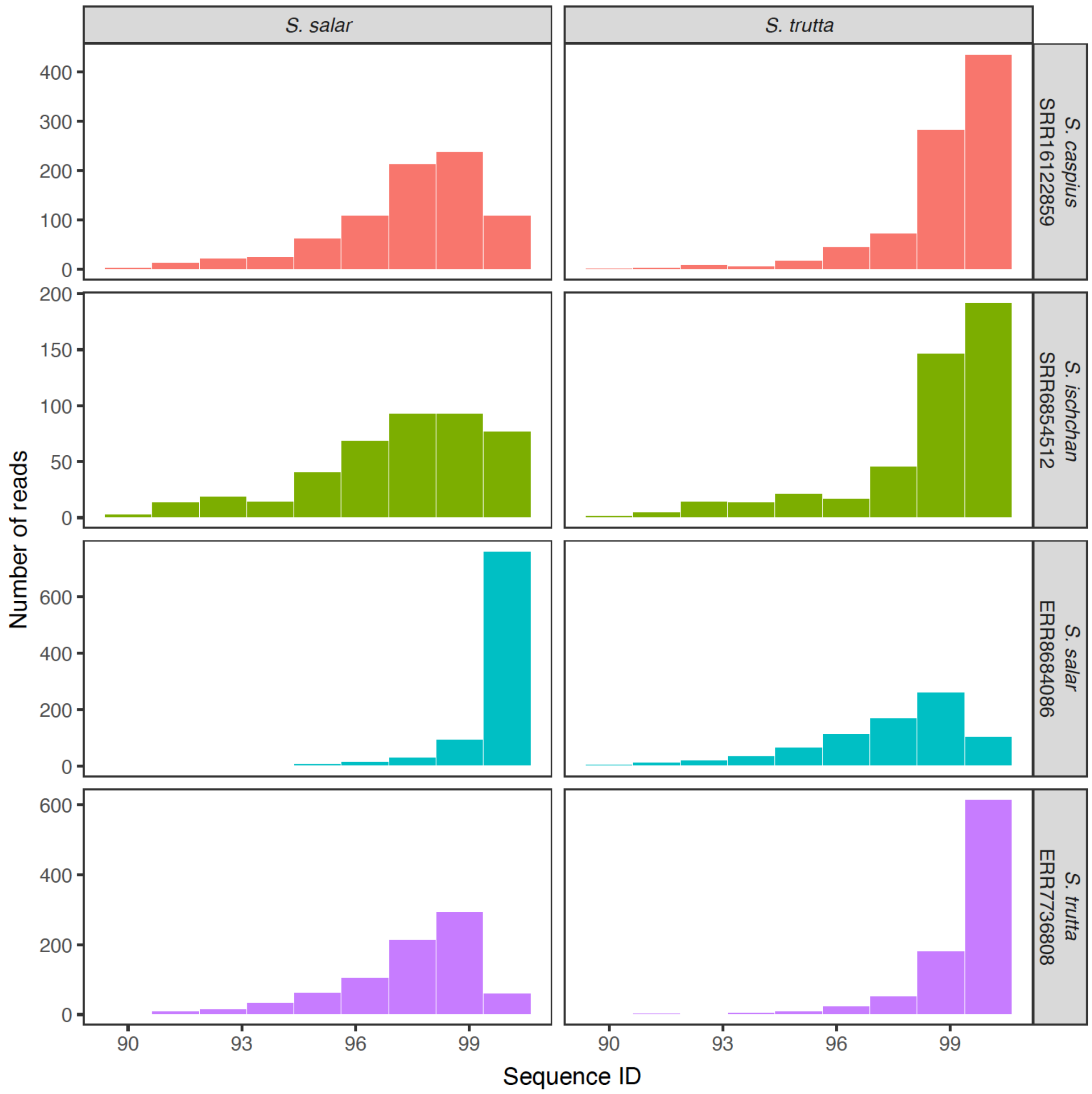


**Figure S1.6.** Intra- and interspecific identity distributions of 1000 shotgun whole-genome sequences of two *Salmo* species. BLAST searches were performed against four congeneric taxa (accession numbers shown in rightmost column). Intraspecific comparisons generate distributions centered at >99.5% sequence identity and highly left-skewed. Interspecific distributions are centered at lower identities. *Salmo salar* is clearly distinguishable from the other species, namely *S. trutta*. All species with >99.5% sequence identity in interspecific comparisons are part of the brown trout complex and their taxonomic status has been debated.

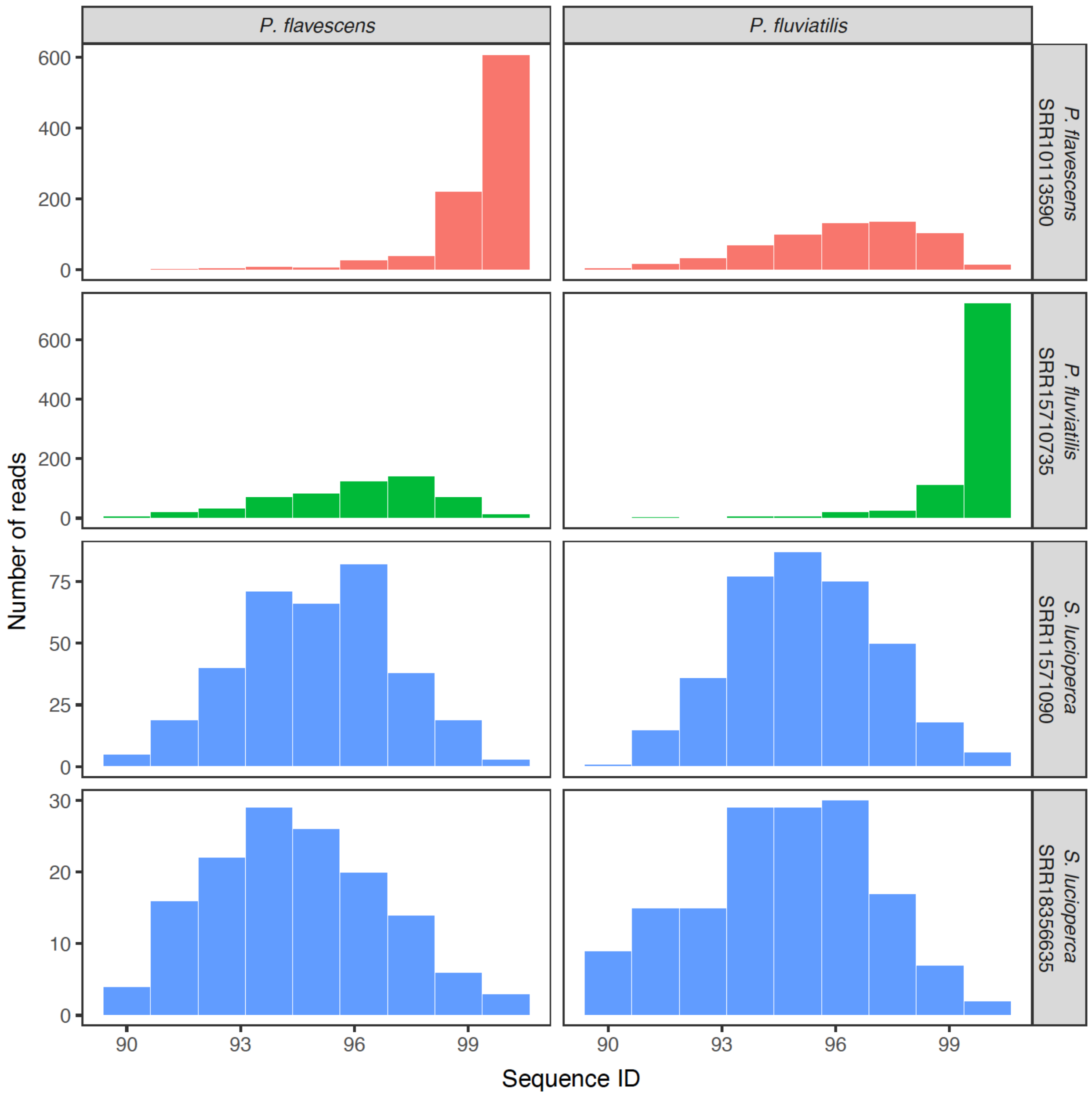


**Figure S1.7.** Intra- and interspecific identity distributions of 1000 shotgun whole-genome sequences of two *Perca* species. BLAST searches were performed against three congeneric taxa (accession numbers shown in rightmost column). Intraspecific comparisons generate distributions centered at >99.5% sequence identity and highly left-skewed. Interspecific distributions are centered at lower identities.

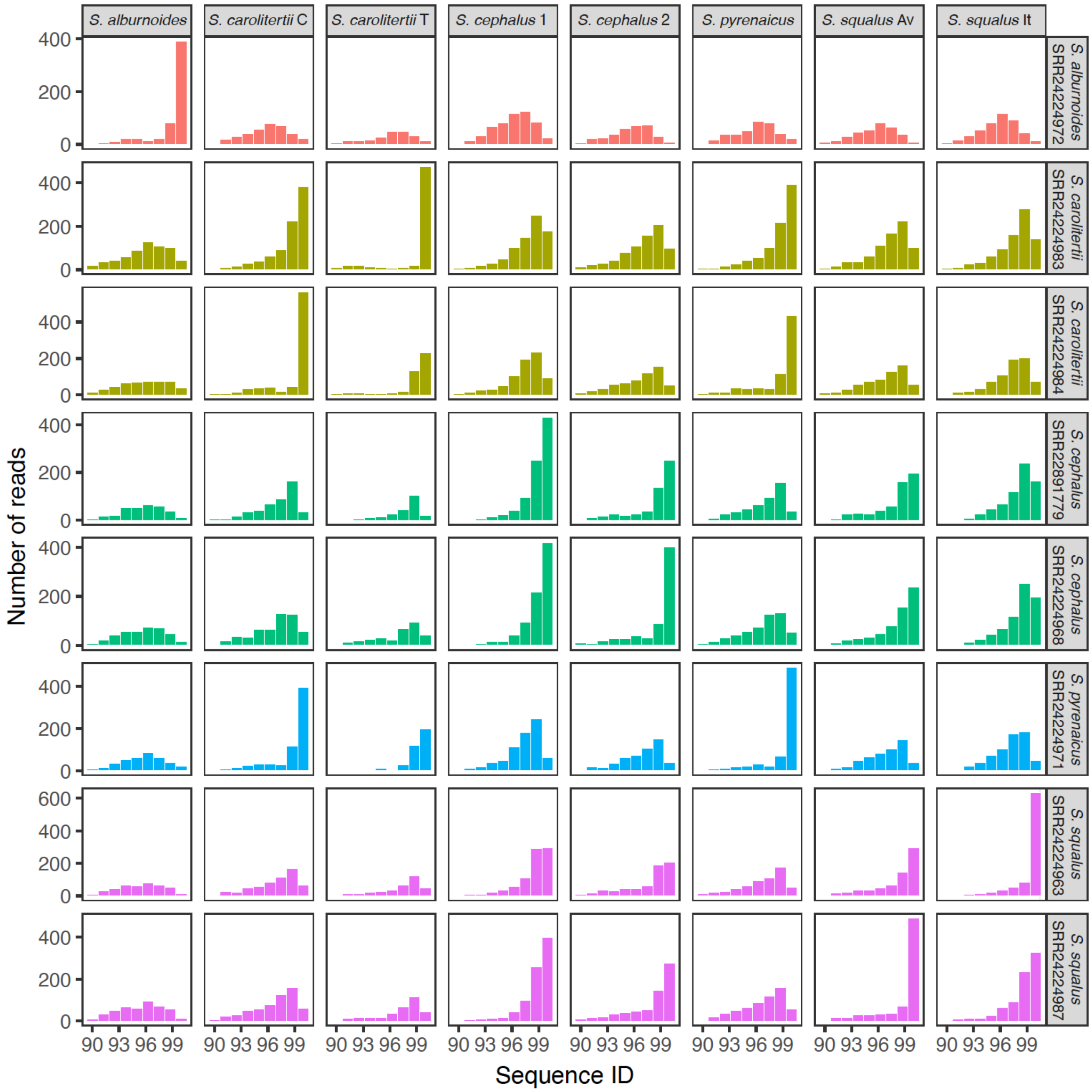


**Figure S1.8.** Intra- and interspecific identity distributions of 1000 shotgun whole-genome sequences of five *Squalius* species. BLAST searches were performed against each other (accession numbers shown in rightmost column). Intraspecific comparisons generate distributions centered at >99.5% sequence identity and highly left-skewed. Interspecific distributions are centered at lower identities.

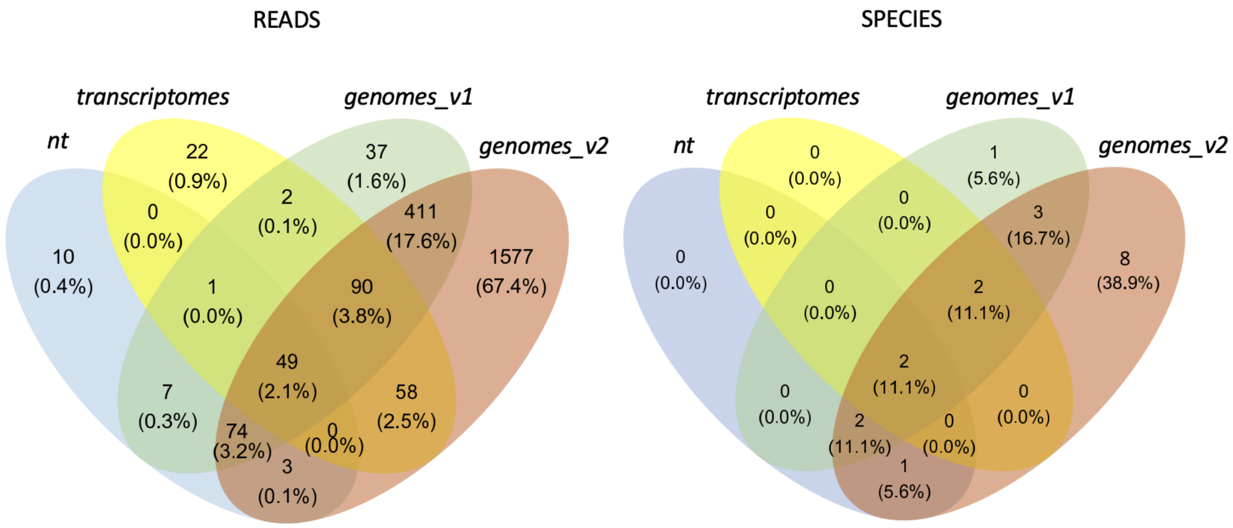


**Figure S2.** Comparison between the four reference databases used to recover matches at the species level for fish taxa. Comparisons are done using the number of reads matching to fish taxa (left) and the number of fish taxa recovered (right). Both *genomes* databases classify more reads and more species, outperforming *nt* and *transcriptomes* databases.

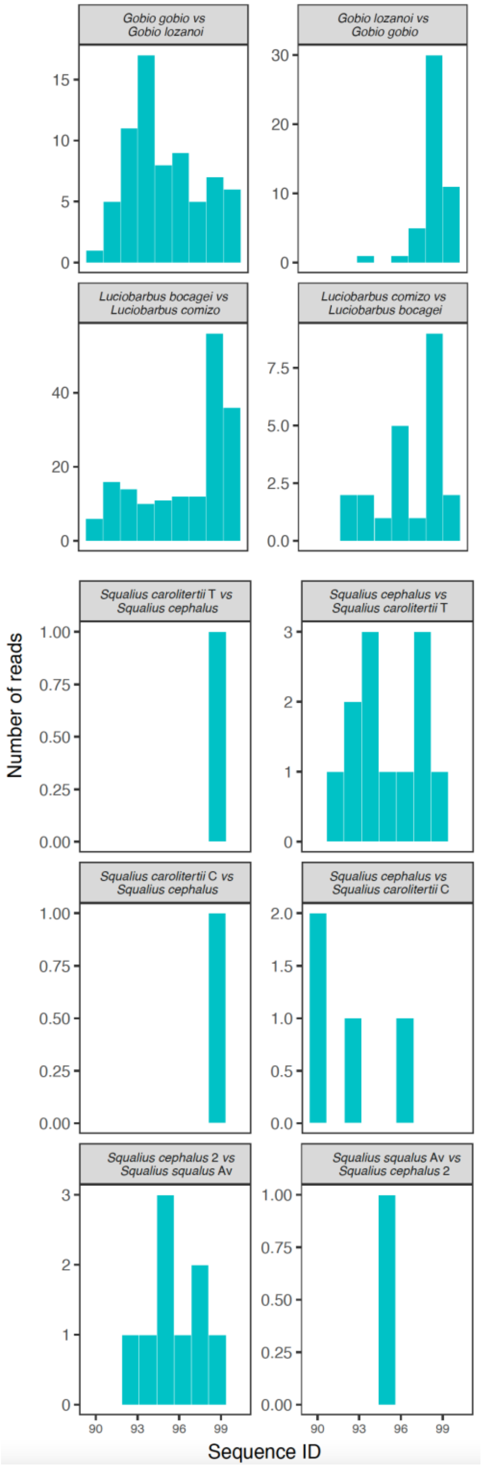


Figure S3. Reciprocal blasts of eDNA reads assigned to five species pairs. BLAST searches were performed by remapping reads against a congeneric species. All interspecific comparisons generate distributions centered at <99.5% sequence identity.

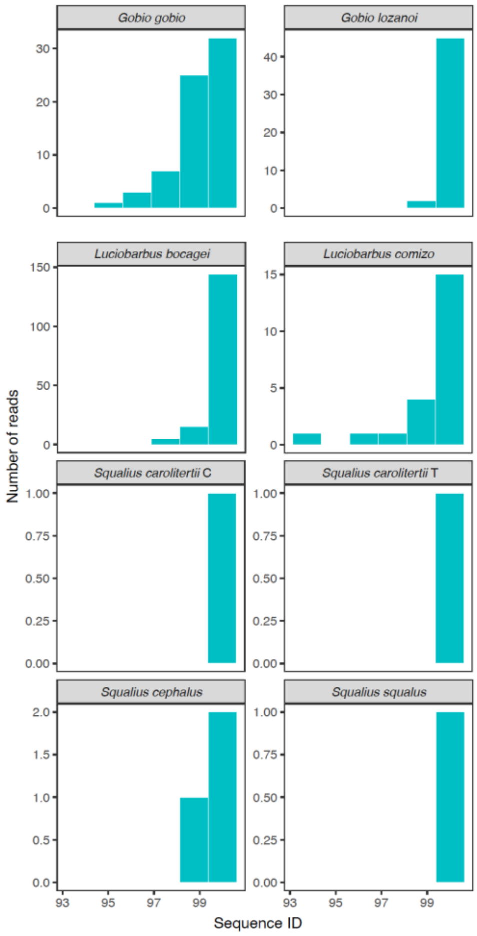


**Figure S4.** Recursive blasts of eDNA reads assigned to five species pairs. BLAST searches were performed by remapping reads back to original species. All generated distributions are centered at >99.5% sequence identity.

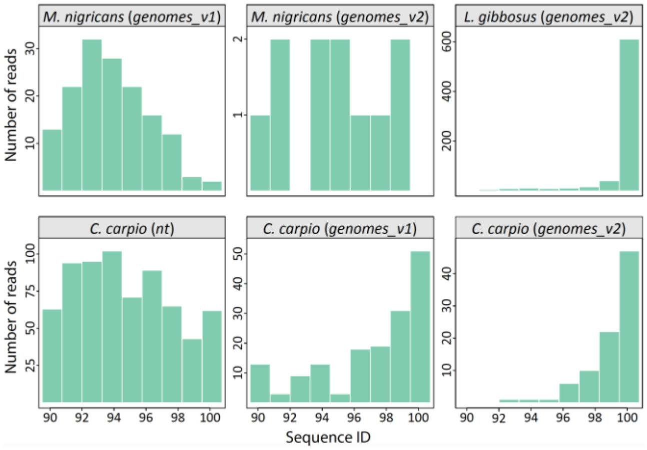


**Figure S5.** Sequence identity distributions of metagenomics reads matching *Micropterus nigricans*, *Lepomis gibbosus* and *Cyprinus carpio* in three different databases: *nt*, fish *genomes_v1* and extended fish *genomes_v2*. The large number of blast hits to *M. nigricans* (*genomes_v1*) and *C. carpio* (*nt* and *genomes_v1*) with low sequence identities are evidence of dark biodiversity, i.e., presence of species not represented in the reference databases. By adding new fish genomes of the suspected dark taxa (*genomes_v2* database), these reads remap to the new genomes with high identity (e.g., *L. gibbosus*), decreasing the number of low-identity matches to *M. nigricans* and *C. carpio*.

### *Supplementary Notes*

#### *Note S1. Criteria to disentangle matches of closely related species*

In the shotgun whole-genome sequence data (sgWGS) comparisons, exceptionally, cases of very closely related species (e.g., *Micropterus* sp., *Squalius* sp. and *Salmo* sp.), in which incomplete lineage sorting (ILS) and introgressive hybridization can contribute to a relatively high proportion of shared genomic regions, sequence identity distributions can be centered at 100% (Figure S1). These false positives might pose challenges when metagenomics is used for *discovery* of new, very closely related taxa, but not in the case of *monitoring* of known taxa. In the latter, all sequence identity distributions can be used in the detection and subsequent exclusion of false positives, increasing the accuracy of the metagenomics assessment. Such criteria were used to disentangle the matches from three genera (*Luciobarbus, Gobio,* and *Squalius*) showing distributions with modes at 100% for two congeners (Figure 3).

Introgression may be one driver of false positives: it has been reported for many of species pairs, such as between *L. bocagei* and *L. comizo,* and between *S. cephalus* and *S. squalus* (Buj et al., 2020; Gante et al., 2015; Segherloo et al., 2021). Thus, it is likely that the specimens used for genome sequencing harbor some genetic signal from the other species, which would result in their molecular detection, i.e., false positives. Previous evidence of introgressive hybridization is strong for *Luciobarbus*, and the identity distributions obtained for the comparison between *L. bocagei* and *L. comizo* using shotgun whole genome sequence data are congruent with an intraspecific pattern instead (Figure S1.5). These are also very young species with long generation times, therefore increasing the chances of ILS. Thus, the taxonomic identification of *Luciobarbus* is conservatively made considering a “composite” taxon including both species under the name *L. bocagei,* since *L. comizo* has never been recorded north of the Tagus River. Given the history of hybridization between these taxa, detection using metabarcoding is also affected and only done at the genus level.

In what concerns the *Squalius* species pair mentioned above, we determine that the individuals originating from the Ave River are themselves interspecific hybrids, as their genomes show high sequence identity to both *S. squalus* and *S. cephalus* as opposed to another *S. squalus* sample originating from its native distribution in Italy (Figure S1.8). Preliminary morphological data on collected specimens also indicates intermediacy between the two species. Thus, while the two genetic signals detected are in fact real, they originate from one admixed biological entity. This level of resolution is only possible with our multilocus analyses of metagenomics data, as metabarcoding only detects one mitochondrial type (*S. squalus*). Hence, we exclude *S. cephalus* from the list of species present in the Ave River.

Finally, we consider the first molecular detection of *G. gobio* in Portugal as a true positive, considering the intra- and interspecific distributions (Figure S1.3). Whether *G. gobio* specimens are locally present in the Ave River, or this species has introgressed the Ave River *G. lozanoi* population warrants further research, as only one mitochondrial type (*G. lozanoi*) is detected by metabarcoding.

While a species list could conservatively include all taxa in a species discovery scenario, pre-existing information from historical sample collections points to which species from each of the pairs is likely present in the Ave River, minimizing the risk of false positives and resulting in the final 16 taxa detected using multilocus metagenomics taxon assignment.

#### *Note S2. Probability of detection of metabarcoding vs metagenomics*

The probability of a specific locus being sampled in a metagenomics assay is given by the following formula described in (Beszteri et al., 2010): $p_m \cong l_m/G \cdot C_m$; where $p_m$ is the probability of sampling the locus $m$, $l_m$ is the number of nucleotides covered by locus $m$, $G$ the genome size and $C_m$ is the copy number of the locus $m$. Considering the turquoise killifish (*Nothobranchius furzeri*), whose maximum mitochondria count per cell is around 1000 (Hartmann et al., 2011) and whose haploid genome size is at least 1.6 Gbp (Reichwald et al., 2009), all else being equal, the probability of sampling the MiFish marker ($l_m$ = 180 bp; $C_m$ = 1000; and $G = 1.6 \times 10^9$) would be 18,000 times lower than any nuclear genomic region. This discrepancy in sensitivity due to sampling probability is more relevant in rarer species, especially those with larger nuclear genomes and/or fewer mitochondria.
